# The enteric pathogen *Cryptosporidium parvum* exports proteins into the cytoplasm of the infected host cell

**DOI:** 10.1101/2021.06.04.447155

**Authors:** Jennifer E. Dumaine, Adam Sateriale, Alexis R. Gibson, Amita G. Reddy, Jodi A. Gullicksrud, Emma N. Hunter, Joseph T. Clark, Boris Striepen

## Abstract

The parasite *Cryptosporidium* is responsible for diarrheal disease in young children causing death, malnutrition, and growth delay. *Cryptosporidium* invades enterocytes where it develops in a unique intracellular niche. Infected cells exhibit profound changes in morphology, physiology and transcriptional activity. How the parasite effects these changes is poorly understood. We explored the localization of highly polymorphic proteins and found members of the *C. parvum* MEDLE protein family to be translocated into the cytoplasm of infected cells. All intracellular life stages engage in this export, which occurs after completion of invasion. Mutational studies defined an N-terminal host-targeting motif and demonstrated proteolytic processing at a specific leucine residue. Direct expression of MEDLE2 in mammalian cells triggered an ER stress response that was also observed during infection. Taken together, our studies reveal the presence of a *Cryptosporidium* secretion system capable of delivering pathogenesis factors into the infected enterocyte.

## INTRODUCTION

The Apicomplexan parasite *Cryptosporidium* is a leading cause of diarrheal disease worldwide. Young children are highly susceptible to infection and cryptosporidiosis is an important contributor to child mortality (Khalil et al., 2018; Kotloff et al., 2013). Children in resource poor settings carry a disproportionate burden of severe disease (Choy and Huston, 2020). Malnutrition enhances the risk of severe cryptosporidiosis, and at the same time, the disease impacts the nutritional state of children, which can lead to impaired growth (Costa et al., 2011; Mondal et al., 2009). Infection with the parasite results in protective immunity, but this immunity is not sterile and may require multiple exposures to develop (Chappell et al., 1999; Okhuysen et al., 1998). Most human disease is due to infection with *C. hominis,* which only infects humans, and *C. parvum,* which can be zoonotically transmitted (Feng et al., 2018; Nader et al., 2019). The emergence of *Cryptosporidium* species is driven by host adaptation resulting in specialization and narrowing host specificity; however, the sexual lifecycle of the parasite allows for recombination and can lead to rapid convergent evolution of host specificity (Guo et al., 2015; Nader et al., 2019).

*Cryptosporidium* infects the epithelium of the small intestine, where it lives in a unique intracellular, but extracytoplasmic niche (Elliott and Clark, 2000). The mechanism by which this niche is established during invasion is still debated but involves the rearrangement of the host actin cytoskeleton, the formation of tight junction-like structures between host and parasite membranes, and a dense band of unknown composition at the host-parasite interphase (Bonnin et al., 1999; Elliott and Clark, 2000). *Cryptosporidium* has severely reduced metabolic capabilities and relies heavily upon the host cell for nutrients and metabolites (Abrahamsen et al., 2004; Xu et al., 2004). A number of specialized uptake mechanisms have been proposed to fill this need, many of which are believed to be localized to the so-called feeder organelle at the host parasite interface (Perkins et al., 1999). In summary, *Cryptosporidium* remodels the host cell in significant ways that include its cytoskeleton (Bonnin et al., 1999; Elliott and Clark, 2000), cellular physiology and metabolism (Argenzio et al., 1990; Kumar et al., 2018), as well as aspects of immune restriction and regulation (Laurent and Lacroix-Lamandé, 2017).

Many bacterial, protozoan and fungal pathogens use translocated effectors to manipulate their hosts to secure nutrients and to block host immunity. In *Plasmodium falciparum*, exported effectors form adhesive structures on the surface of red blood cells to alter tissue distribution and mechanical properties to prevent clearance (Crabb et al., 1997; Leech et al., 1984) and install new nutrient and ion uptake mechanisms (Baumeister et al., 2006). In *Toxoplasma gondii*, translocated effectors disarm critical elements of interferon induced cellular restriction (Gay et al., 2016; Olias et al., 2016), establish access to cellular nutrients, and rewire signaling and transcriptional networks to antagonize immune responses to promote a favorable environment for parasite growth (Bougdour et al., 2013; Braun et al., 2013; Braun et al., 2019; Gold et al., 2015). *Cryptosporidium* has been hypothesized to use exported effector proteins to ensure its survival (Pellé et al., 2015); however, to date, no such factors have been identified. Here we report that the polymorphic protein MEDLE2 is exported to the host cell cytoplasm by all stages of the *C. parvum* lifecycle *in vitro* and *in vivo*. This protein is not injected during invasion, but rather is transported into the host cell following the initial establishment of infection. We carefully mapped the requirements for export and found a signal that includes a proteolytic cleavage site, and we demonstrate that exported proteins undergo processing. Overall, we demonstrate the presence of a robust translocation mechanism established by intracellular parasites that delivers parasite pathogenesis factors to the host cell.

## RESULTS

### The *Cryptosporidium parvum* protein MEDLE2 is exported into the host cell

The genome of *Cryptosporidium parvum* encodes multiple families of paralogous proteins which carry N-terminal signal peptides and share conserved amino acid repeat motifs among family members (Abrahamsen et al., 2004). These genes are often found within clusters (homologous and heterologous), many of which are located proximal to the telomeres of multiple chromosomes (Figure 1A). Comparing strains and species, these genes are highly polymorphic and vary in copy number, which has been interpreted as sign of rapid evolution driven by their roles in invasion, pathogenesis, and host cell specificity (Guo et al., 2015; Nader et al., 2019; Xu et al., 2019) We hypothesized that such roles might be reflected in the targeting of presumptive effectors to the host cell and selected representatives from each polymorphic gene family for initial localization studies (Supplementary Table 1). Selected loci were modified in the *C. parvum* IOWAII isolate using CRISPR/Cas9 driven homologous recombination (Vinayak et al., 2015) to append three hemagglutinin epitopes (3XHA) in translational fusion to the C-terminus (Figure 1A). Drug resistant parasites were recovered for four of six initial candidates and successful genomic insertion was mapped by PCR (Figure 1B and Figure1-figure supplement 1). We next infected human ileocecal colorectal adenocarcinoma cell cultures (HCT-8) with transgenic parasites and assessed the localization of the tagged proteins by immunofluorescence assay (IFA, Figure 1D & E). For most candidates, the tagged protein (red) appeared to coincide with the parasite and/or the parasitophorous vacuole (cgd8_3560, Figure 1D, Figure 1-figure supplement 1). In contrast, upon infection with parasites tagged in the MEDLE2 locus (cgd5_4590), HCT-8 cells showed HA staining in the cytosol (Figure 1E). We note that the cells that stain for HA (red) were those that were infected with parasites, labeled with *Vicia villosa* lectin (VVL, green), and conclude that MEDLE2-HA is exported by the parasite into the host cell during or following invasion.

**Figure 1.**
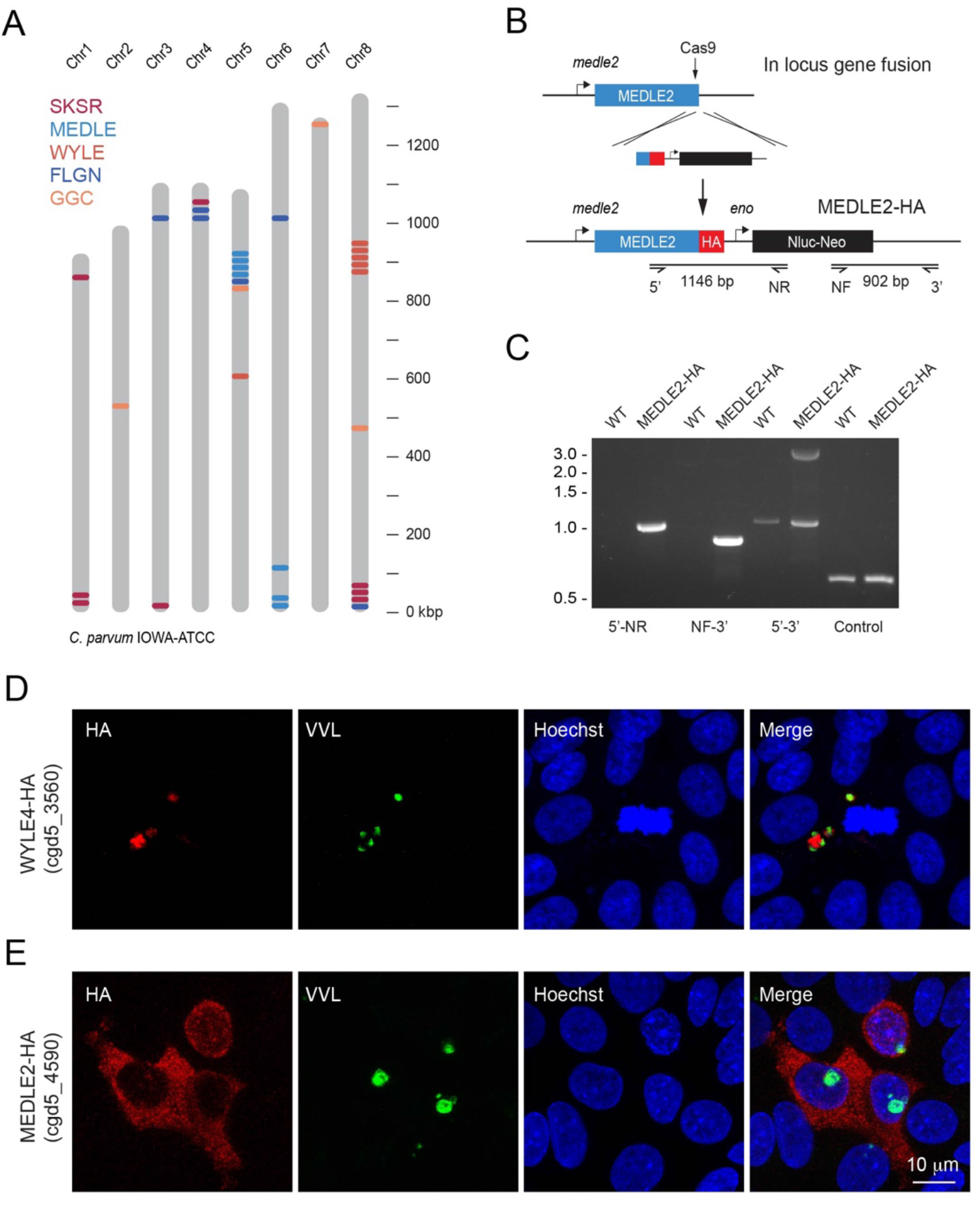
MEDLE2 is exported to the host cell cytoplasm. (A) Schematic overview of the chromosomal location for polymorphic gene families in the *C. parvum* genome. (B) Map of the MEDLE2 locus targeted in *C. parvum* for insertion of a 3X hemagglutinin epitope tag (HA), a nanoluciferase reporter gene (Nluc), and neomycin phosphotransferase selection marker (Neo). (C) PCR mapping of the MEDLE2 locus using genomic DNA from wild type (WT) and transgenic (MEDLE2-HA) sporozoites, corresponding primer pairs are shown in B, and thymidine kinase (TK) gene used as a control. Note, the presence of two bands in the 5’ – 3’ amplification, indicating the presence of a transgene (3081 bp) and persistence of an unmodified copy (1174 bp), suggesting multiple copies of MEDLE2 in the *C. parvum* genome, also see Figure 1-figure supplement 2. (D-E) HCT-8 cultures were infected with WYLE4-HA (D) or MEDLE2-HA (E) transgenic parasites and fixed after 24h for IFA. Red, antibody to HA; green, *Vicia villosa* lectin stain, VVL (Gut and Nelson, 1999); blue, Hoechst DNA dye. Additional genes targeted and the localizations of their products are summarized in Supplemental Table 1 and Figure1-figure supplement 1 and Figure 1-figure supplement 3

We investigated whether other members of the MEDLE gene family are similarly exported and selected MEDLE1 (cgd5_4580) and MEDLE6 (cgd6_5490) for epitope tagging. In IFAs, both tagged MEDLE1 and MEDLE6 localized around the parasite, as well as to the host cell cytoplasm, but expression and export was less than what was observed for MEDLE2 (Figure 1- figure supplement 3). To explore this difference, we engineered a chimeric mutant carrying an extra copy of MEDLE1-HA driven by the MEDLE2 promoter. Upon infection with these parasites, we observed increased MEDLE1 expression making export into the host cell easier to appreciate (Figure1-figure supplement 3). We conclude that multiple members of the MEDLE family are host targeted proteins. Of these, MEDLE2 was expressed and exported most robustly and was selected for further study.

### MEDLE2 is exported across the *C. parvum* lifecycle in culture and infected animals

To better understand export of MEDLE2 to the host cell, we next engineered a reporter parasite in which the endogenous locus of MEDLE2 was 3XHA epitope tagged, and these parasites expressed a tandem mNeon green fluorescent protein in their cytoplasm to allow for identification and quantification of parasites (MEDLE2-HA-tdNeon, Figure 2- figure supplement 1). These parasites were then used in time course experiments across the 72 hours of infection afforded by the HCT-8 culture model. Following inoculation with sporozoites, cultures were fixed in 12-hour increments and processed for IFA. MEDLE2 was observed at all timepoints (Figure 2A) and quantification showed that the number of HA positive host cells (red, Figure 2B) increased over time, closely matching the increase in the number of parasites (blue). 94 % ± 1.83 (Mean ± SD, *n*= 3695) of the cells showing HA staining also showed parasite infection. Importantly, this high level of correlation between host cell HA expression and infection (r^2^ = 0.9) remained constant over 72 hours. Previous work has shown synchronous lifecycle progression over this time span. Initially, all parasites represent asexual stages replicating by merogony followed by a dramatic development switch at 40 hours, and later cultures are dominated by male and female gametes (Tandel et al., 2019). We staged parasites at 48 hours and identified female gametes and male gamonts using antibodies for the markers COWP1 and tubulin, respectively. We found the host cells infected with all of these stages to be positive for MEDLE2-HA (Figure 2D).

**Figure 2.**
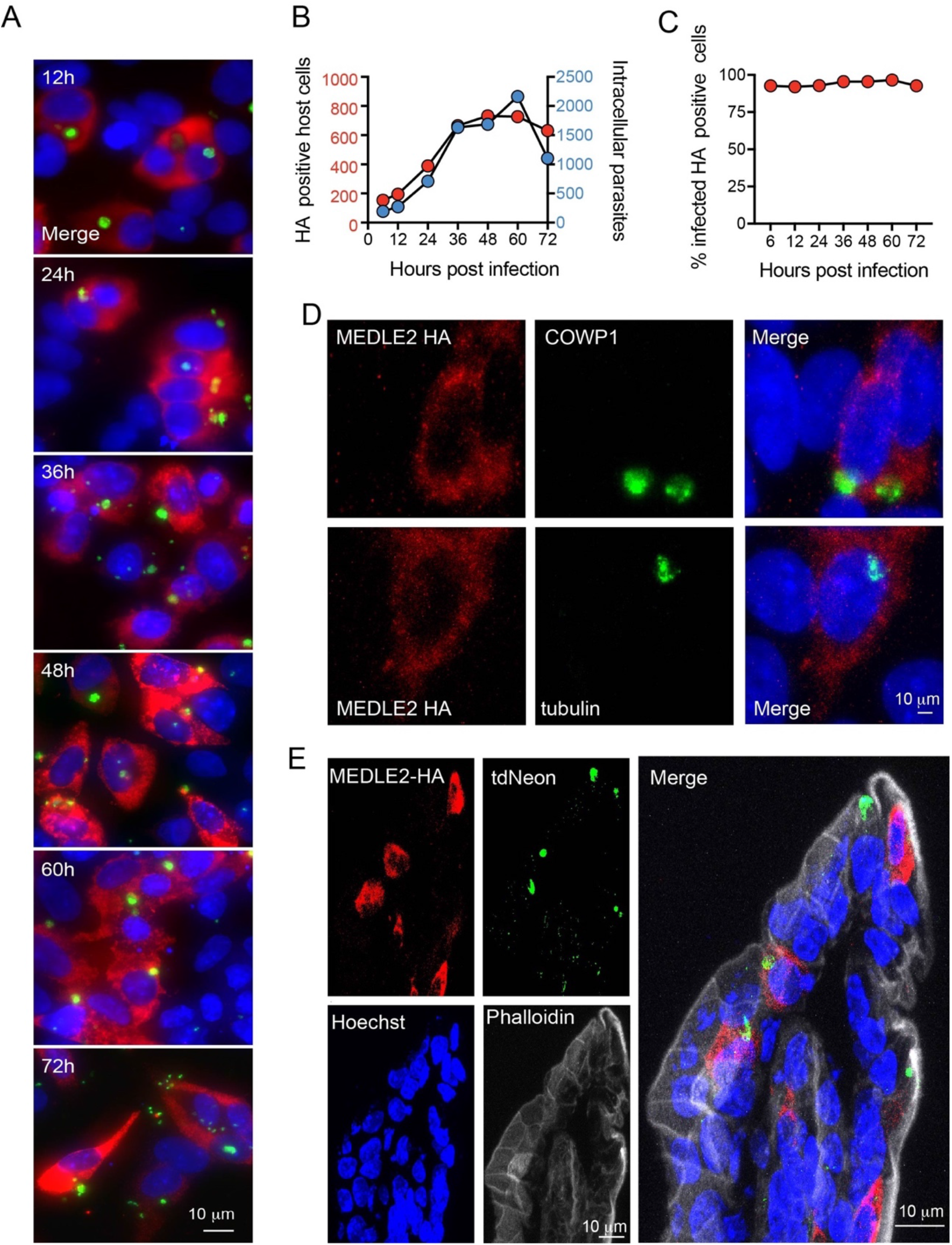
Infected cells express MEDLE2-HA across the parasite lifecycle. (A) 200,000 MEDLE2-HA-tdNeon transgenic parasites were used to infect HCT8 cells and fixed at intervals across a 72h time period. Data shown are representative images from triplicate coverslips processed for IFA. Red, HA tagged protein; green, parasites (mNeon); blue, Hoechst. (B-C) Quantification of MEDLE2 expressing cells (red) versus intracellular parasites (blue) for 3695 host cells evaluated across a 72h time course. 20 fields of view quantified using ImageJ to identify host cells and parasites (B). The percentage of cell exhibiting MEDLE2-HA and mNeon staining is constant across the time course with a cumulative 94 % ± 1.83 (Mean ± SD) (C). (D) HCT-8 cultures infected with MEDLE2-HA parasites were fixed for IFA at 48h when sexual life stages were present. Cells were stained with stage specific antibodies for female (COWP1) and male (α-tubulin) demonstrating MEDLE2 is exported across the parasite lifecycle. Red, HA tagged protein; green, parasites (stage specific antibody); blue, Hoechst. (E) IFA of cryosections from the small intestine of *Ifng^-/-^* mice infected with MEDLE2-HA-tdNeon *C. parvum* (images representative of samples from 3 mice). Red, HA tagged protein; green, parasites (tdNeon); blue, Hoechst; grey, Phalloidin (actin).

We also tested whether MEDLE2 export occurs *in vivo*. Susceptible *Ifng^-/-^* mice were infected with 10,000 oocysts of the reporter strain, and after 12 days, mice were euthanized, the ileum was resected, fixed, and processed for histology. As shown by immunohistochemistry of sections of infected intestines in Figure 2E, MEDLE2 was exported to infected cells *in vivo* and exhibited cytoplasmic localization.

### MEDLE2 is expressed and exported by trophozoites once infection has been established

Two broadly conserved temporal patterns have been described for host-targeted effectors in Apicomplexa (Figure 3A). Those involved in early aspects of the infection are packaged into the rhoptry organelle and injected during invasion (Rastogi et al., 2019). A second wave of proteins is translated and exported after the parasite has established its intracellular niche and they are delivered to the host cell by a translocon-based mechanism (de Koning-Ward et al., 2009; Franco et al., 2016). For this reason, we next determined the timing of MEDLE2 expression using IFA of wild type (WT) and transgenic sporozoites mounted to cover glass with poly-lysine. We readily observed labeling for the sporozoite antigen Cp23 but did not detect any HA staining in transgenic sporozoites, indicating that MEDLE2 is not pre-packaged into secretory organelles (Figure 3B). We then stained intracellular stages at different timepoints following invasion to determine the kinetics of MEDLE2 expression and export. At 4 hours, HA is first detectable, with labeling being associated with the parasite (parasite nuclei stained with Hoechst highlighted by white arrowheads). Beginning at 5.5 hours, MEDLE2-HA staining was observed throughout the host cell cytoplasm and continued to accumulate over time (Figure 3C).

**Figure 3.**
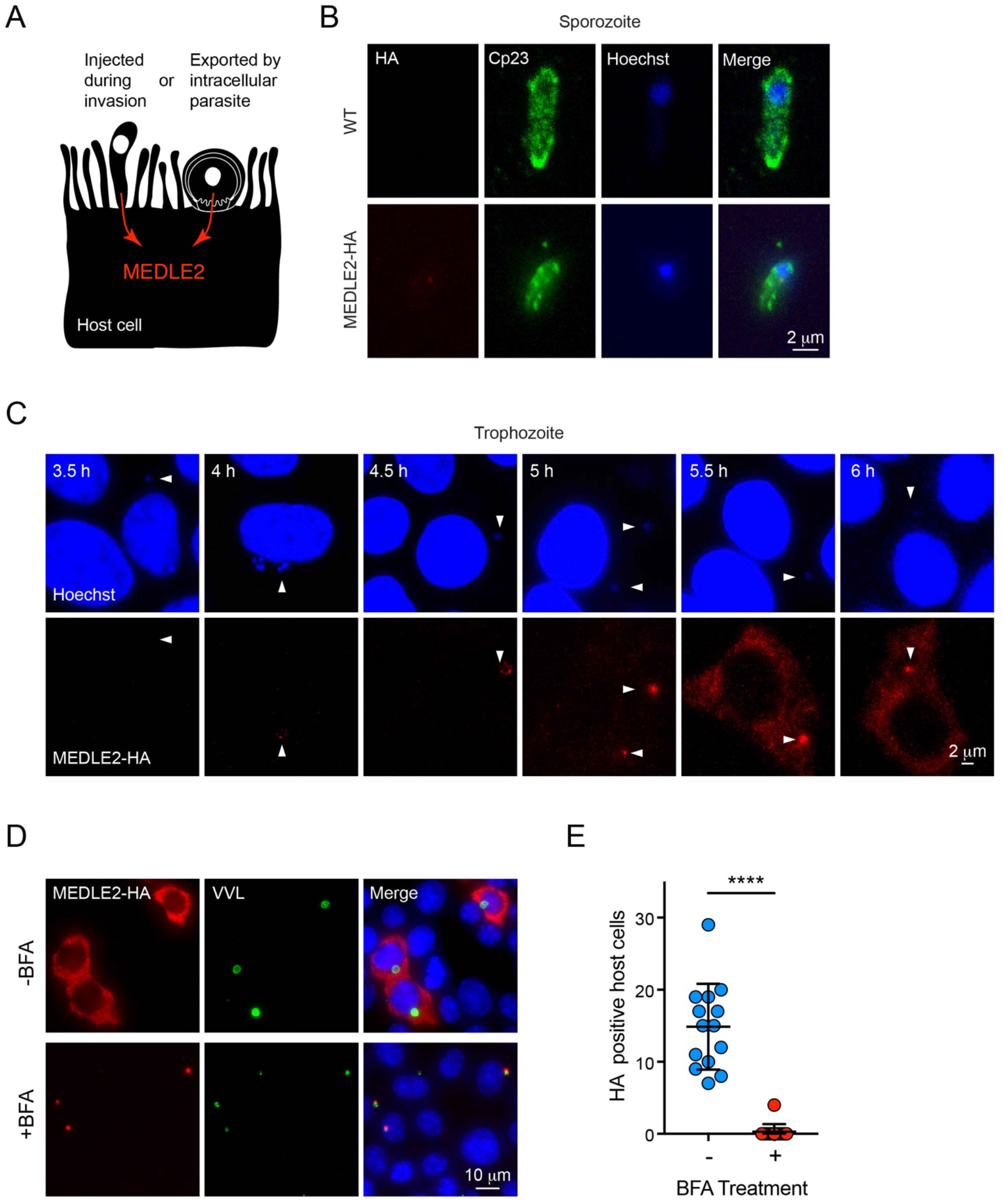
MEDLE2 is expressed by trophozoites and passes through the secretory pathway. (A) Schematic representation of hypothetical patterns of MEDLE2 export in *C. parvum*. (B) IFA of WT and MEDLE2-HA sporozoites fixed on Poly-L-Lysine treated coverslips. We note that MEDLE2-HA is not observed in sporozoites. Red, HA tagged protein; green, sporozoite antigen Cp23; blue, Hoechst. (C) HCT-8 cells infected with MEDLE2-HA parasites were fixed in 30 min increments and processed for IFA. Data shown are representative images from a time course bridging 3h (no observed MEDLE2-HA) and 6h (MEDLE2-HA abundant in host cell). White arrowheads denote parasite nuclei. Red, HA tagged protein; blue, Hoechst. (D) MEDLE2-HA parasites were excysted and used to infect HCT8 and after 3h media were supplemented with BFA (10 ug/mL). 10h post infection, cells were fixed and processed for IFA. Red, HA tagged protein; green, parasites (VVL); blue, Hoechst (D). (E) The impact of BFA treatment on MEDLE2-HA export was quantified showing a significant reduction in MEDLE2 export when comparing BFA treated (red) and untreated cells (blue), (*n =* 191 untreated, *n =* 98 treated; mean ± SEM; *p* < 0.0001; unpaired *t* test with Welch’s correction).

MEDLE2 is predicted to be a 209 amino acid protein with a putative N-terminal signal peptide suggesting trafficking through the parasite’s secretory pathway. To test this, we used Brefeldin A (BFA) which blocks passage through the secretory pathway between the ER and Golgi. Cells were infected with MEDLE2-HA parasites, treated with BFA beginning 3 hours post-infection, and fixed for IFA after 10 hours to assess MEDLE2 localization. BFA treatment initiated after invasion ablated export and resulted in the accumulation of MEDLE2-HA within the parasite (Figure 3D). Image analysis and quantification showed this reduction in export to be significant when comparing treated (red) to untreated cells (blue) (*n* = 98 treated; *n =* 198 untreated*; p* < 0.0001; unpaired *t* test with Welch’s correction; Figure 3E). We conclude that MEDLE2 is not injected by the sporozoite during invasion; rather, it is expressed and exported by the trophozoite and arrives in the host cell about 5 hours post infection (note that the length of the intracellular lytic cycle of asexual stages is 11.5 hours (Guerin & Striepen unpublished).

### MEDLE2 is an intrinsically disordered protein and its export is blocked by ordered reporters

To further investigate the export of MEDLE2, we sought to develop a reporter assay to follow trafficking using three previously established systems. We fused the fluorescent reporter mScarlet to the C-terminus of MEDLE2 in its endogenous locus (Figure 4A and Figure 4-figure supplement 1). Transgenic parasites showed robust expression of the reporter when used to infect HCT-8 cultures. However, this fluorescence (red) was associated with parasites, highlighted by staining with VVL (green, Figure 4B), suggesting that MEDLE2-mScarlet remained trapped within or close to the parasite. We considered that export may occur, but that we lack the sensitivity to detect it. Thus, we engineered two highly sensitive assays that employ enzymes to amplify the signal.

**Figure 4.**
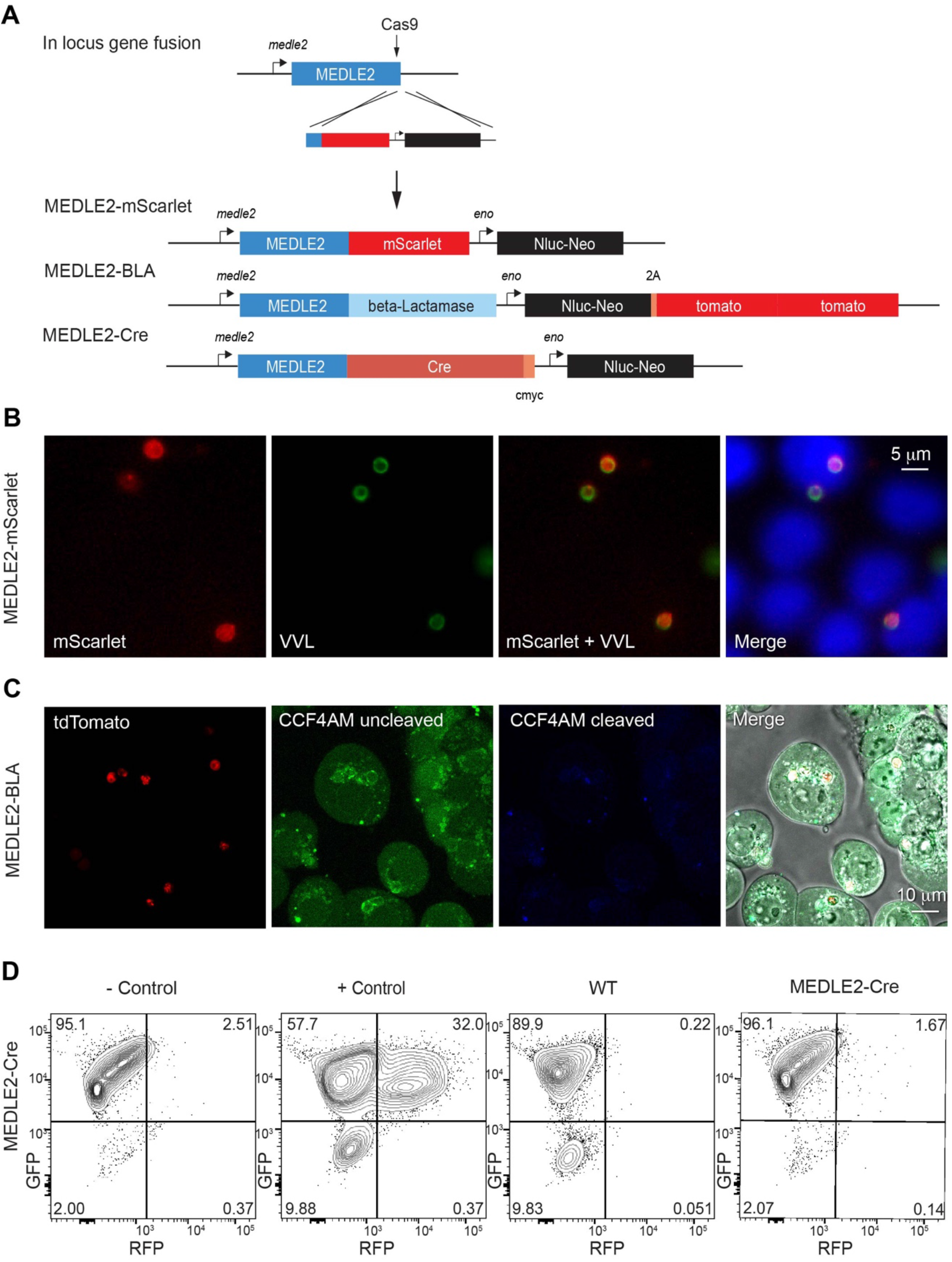
Ordered reporters disrupt MEDLE2 export. (A) Schematic map of the MEDLE2 locus targeted for insertion of three different reporter genes (mScarlet, Beta-lactamase, or Cre recombinase), nanoluciferase (Nluc), and the selection marker (Neo). The guide RNA and flanking sequences used here were the same as those employed to generate MEDLE2-HA transgenic parasites (see Figure 2-figure supplement 1, Figure 4-figure supplement 1 for more detail). (B) MEDLE2-mScarlet parasites were used to infect HCT8 cells and fixed for IFA across a time course. Data shown are from 10h post-infection, which is representative of the MEDLE2 localization observed at all time points. Red, Medle2-mScarlet; green, parasites (VVL); blue, Hoechst. (C) HCT8 cells were infected with MEDLE2-BLA *C. parvum* for 24h before incubation with the CCF4-AM Beta-lactamase substrate and visualization by live microscopy. This experiment was repeated 3 times. Red, parasites (tdTomato); green, uncleaved CC4F-AM; blue, cleaved CCF4-AM; grey, DIC. We attribute lack of CCF4-AM cleavage to failure of MEDLE2-BLA to export (Figure 4-figure supplement 1). (D) MEDLE2-Cre parasites were used to infect loxGFP/RFP color switch HCT8 cells (Figure4- figure supplement 2) for schematic representation). After 48h, cells were subjected to flow cytometry. Live, single cells were gated based upon forward and side scatter, and green fluorescence (GFP) and red fluorescence (RFP) were measured to detect Cre recombinase activity. Despite robust infection, MEDLE2-Cre infected cultures did not express RFP (Figure 4- figure supplement 2), compared to the positive control which was transiently transfected to express Cre recombinase.

First, we tagged MEDLE2 with beta lactamase, which has been used to reveal effector export in bacteria and protozoa (Charpentier and Oswald, 2004; Lodoen et al., 2010). These parasites also expressed a cytoplasmic red fluorescent protein (Figure 4-figure supplement 1B). Cells infected with MEDLE2-BLA sporozoites were incubated with CCF4-AM, a cell-permeable substrate of beta lactamase and imaged by live microscopy. Infected and uninfected cells accumulated CCF4-AM (green, Figure 4C); however, we did not detect substrate cleavage resulting in blue fluorescence (Figure 4C). This could be due to the lack of MEDLE-BLA expression or export. To visualize localization of the MEDLE2-BLA fusion protein during infection, we performed IFA on MEDLE2-BLA infected cells using a BLA antibody and observed that MEDLE2-BLA (green) was expressed but remained with the parasite (red, Figure 4-figure supplement 1C). Next, we generated a MEDLE2 fusion with Cre recombinase and used these parasites to infect an HCT-8 host cell line we engineered to carry a floxed GFP/RFP color switch reporter (see Figure4- Figure Supplement 2pdf). After 48 hours, cells were subjected to flow cytometry (Figure 4D). Using a gate for live, single cells, uninfected cells showed green (95.1%) but not red fluorescence (2.51%). Transfection of host cells with a Cre recombinase expression plasmid resulted in a pronounced shift to red fluorescence (+control, 32%, Figure 4D). Cells infected with either WT parasites or MEDLE2-Cre parasites remained green, despite robust infection (Figure 4-figure supplement 2). Therefore, we tested three reporters, none of which resulted in detectable export to the host cell. We note that multiple algorithms predict MEDLE2 to be a highly disordered protein (low complexity regions are indicated in light blue in Figure 5B) and conclude that translocation is blocked when folded reporters are fused to the protein.

**Figure 5.**
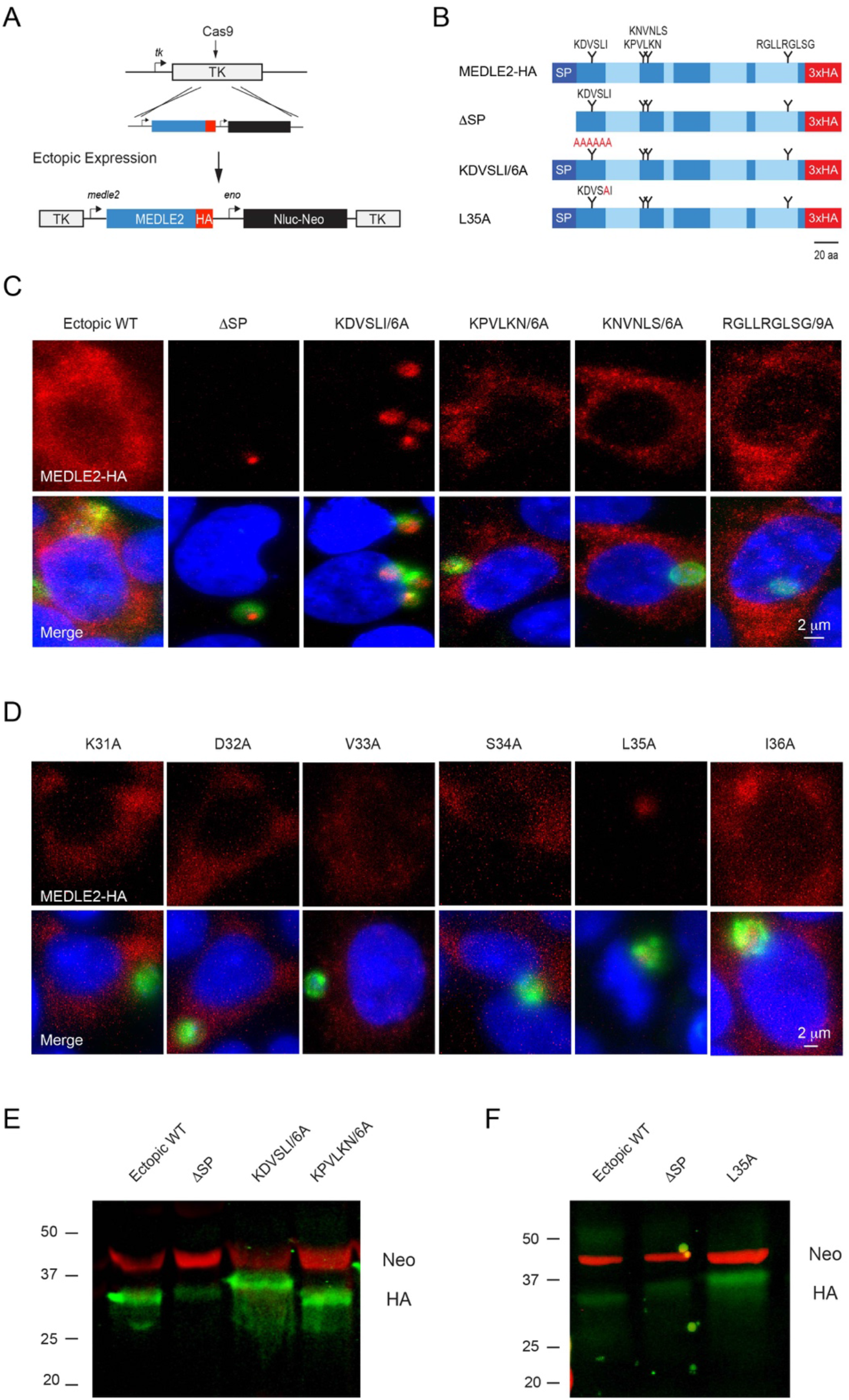
MEDLE2 contains a host targeting motif that is processed during export. (A) Map showing the strategy used to engineer an ectopic copy of MEDLE2-HA in the thymidine kinase (TK) locus. Expression of an ectopic copy of MEDLE2-HA was driven the MEDLE2 promoter. All point mutations were confirmed by Sanger sequencing (Figure 5-figure supplement 1). (B) Schematic representation of the MEDLE2 mutants generated using the strategy outlined in A. The signal peptide (SP) is represented by dark blue and low complexity regions are shown in light blue. Candidate motifs targeted for mutagenesis are indicated with black triangles and mutagenized amino acids are shown in red for two representative mutants. (C-D) Mutant parasites were used to infect HCT8 cells and fixed for IFA after 24h. For mutants shown in (C) the entire candidate motif was replaced with a matching number of alanine residues (ex: KDVSLI/6A à AAAAAA). For mutants shown in (D) each individual amino acid in the KDVSLI sequence was changed to alanine. Red, HA tagged protein; green, parasites (VVL); blue, Hoechst. We note that signal peptide and leucine 35 within the KDVSLI sequence are required for MEDLE2 export. (E-F) 5 × 10^6^ transgenic oocysts were used to infect HCT8 cells for 48h before preparation of whole cell lysates. Proteins were separated by for SDS PAGE and analyzed by Western Blot. The resulting blots for infections with whole motif mutants (E) and individual amino acid point mutants (F) are shown. Red, Neomycin; green, HA. Note that when mutants are expressed in mammalian cells and not C. parvum the resulting proteins do not show any size differences (Figure 5- figure supplement 2).

### MEDLE2 export depends upon N-terminal sequence features

We then sought to determine whether MEDLE2 contains sequence specific information for host-targeting. Using previously established host targeting motifs from *P. falciparum* and *T. gondii* as models (Coffey et al., 2015; Marti et al., 2004), we searched the MEDLE2 amino acid sequence to identify candidate export motifs. Preference was given to regions with a basic amino acid, followed by one or two random amino acids, and a leucine residue (Pellé et al., 2015). While *Plasmodium* host targeting motifs are typically found in close proximity to the signal peptide, *T. gondii* exhibits less rigid distance requirements (Coffey et al., 2015). We identified 4 motifs, three sites in proximity to the N-terminus and one C terminal candidate for mutational analysis (Figure 5A). As folded reporters are not tolerated, we engineered parasite lines to express an ectopic copy of MEDLE2 marked by a HA tag. A cassette driven by the MEDLE2 promoter was inserted into the locus of the dispensable thymidine kinase gene (TK, Figure 5B) and expression level and export of ectopic WT protein was indistinguishable from protein tagged within the native locus.

Removal of the sequence encoding the N-terminal signal peptide (ΔSP) prevented MEDLE2-HA export, and the resulting protein accumulated within the parasite (Figure 5C). Next, we constructed a series of parasite strains in which each of the candidate motifs was replaced by a matching number of alanine residues (all mutants were confirmed by PCR mapping and Sanger sequencing, Figure Figure 5-figure supplement 1). Mutagenesis of three of these candidate motifs had no impact on MEDLE2 translocation to the host cell (Figure 5C; Figure 5- figure supplement 1). In contrast, when the most N-terminal sequence KDVSLI was changed to six alanines, HA staining accumulated in the parasite and host cell staining was lost (Figure 5C). We conclude that in this mutant, MEDLE2-HA is made but export is ablated. Next, we constructed six additional strains using the same strategy to change each amino acid position of the KDVSLI motif to alanine one residue at a time (Figure5-figure supplement 1). Mutation of residue leucine 35 to alanine (L35A) ablated export and instead, MEDLE2-HA remained with the parasite (Figure 5D). Changing the remaining five amino acids individually did not alter MEDLE2 localization in the host cell (Figure 5D). We conclude leucine 35 to be critical for export.

### MEDLE2 export is linked to proteolytic processing

For both *P. falciparum* and *T. gondii*, leucine residues serve as crucial sites for proteolytic processing events that license proteins to leave the parasite and enter the host cell (Coffey et al., 2015; Hiller et al., 2004; Marti et al., 2004). To test whether such processing occurs during export of MEDLE2 in *C. parvum*, we performed Western Blot analysis on whole cell lysates of infected cell cultures using antibodies to HA (MEDLE2-HA, green) and the drug resistance marker Neo (loading control, red). For WT MEDLE2-HA we observed a single band with an apparent molecular weight of 31 kDa (Figure 5E, the predicted molecular weight for full length MEDLE2-HA is 26.9 kDA but the abundance of positive charges is likely to result in reduced electrophoretic mobility). Protein KPVLKN/6A, carrying a mutation that did not affect trafficking to the host cell cytoplasm, was of identical size to WT MEDLE2-HA. In contrast, the KDVSLI/6A mutant, which is no longer exported, appeared to be of a larger molecular weight (Figure 5E). The mutant lacking the 22 amino acid signal peptide (ΔSP) produced an intermediate size band larger than the exported WT but smaller than the retained ΔKDVSLI mutant. We found a very similar pattern when analyzing the single amino acid mutants, where the L35A change caused the mutant protein to migrate more slowly when compared to WT or the ΔSP mutant (Figure 5F). To ensure the observed differences in apparent molecular weight were due to processing by the parasite and not the consequence of folding or subsequent host processing, we also expressed WT and mutants in mammalian cells (see Material and Methods for detail). In this context, the proteins are the same size (Figure 5-figure supplement 2). Overall, we interpret the relative sizes of the mutated proteins to indicate processing of MEDLE2 at a point beyond the signal peptide, a position that would be consistent with leucine 35. We note that this processing appears to require translocation into the ER, as it does not occur in mutants lacking a signal peptide. Furthermore, the L35A mutation apparently prevented removal of the signal peptide, suggesting that processing at L35 could replace the canonical signal peptidase activity.

### MEDLE2 induces an ER stress response in the host cell

To begin to understand the consequence of MEDLE2 export on the host cell, we expressed the protein in human cells. MEDLE2 omitting the N terminal signal peptide (aa 2-20) was codon optimized for human cell expression, appended to the N-terminus of GFP, and the resulting plasmid introduced into HEK293T cells by lipofectamine transfection. The GFP-only parent plasmid served as control. Transfection with both constructs resulted in cytoplasmatic green fluorescence in roughly 40% of cells 24 hours post transfection (Figure 6A). At this time point, GFP positive cells were enriched by flow cytometry and the resulting populations were subjected to mRNA sequencing (3 biological repeats for each sample, Figure 6B). Differential gene expression analysis revealed 413 upregulated genes and 487 genes with lower transcript abundance in MEDLE2-GFP expressing cells compared to cells expressing GFP alone, with an adjusted *p*-value less than 0.05 (Figure 6C). Gene set enrichment analysis showed upregulation in the response to ER stress, including changes in genes linked to the unfolded protein response (UPR). Genes that are part of the core enrichment of the ER stress response are highlighted (red) in the volcano plot and the most upregulated genes are identified by name (Figure 6D). Genes that were differentially expressed at an adjusted *p*-value less than 0.01 were used to derive a MEDLE2 response signature from the transfected cell dataset (234 genes). Using a published mRNA sequencing dataset generated from *C.* parvum infection of a homeostatic mini-intestine model (Nikolaev et al., 2020), we found enrichment for this MEDLE2 response signature with 51 genes of the MEDLE2 response present, and 22 of them contributing to the core enrichment of this response (Figure 6E). To test whether this ER stress response also occurs during *in vivo* infection, we performed qPCR on ileal segments resected from mice infected with *C. parvum* or those that were uninfected. We measured the RNA abundance for the four genes highlighted in the volcano plot and found three to be upregulated in infected mice compared to uninfected controls (NUPR1, CHAC1, DDIT3, Figure 6F). We conclude that an ER stress response triggered by ectopic expression of MEDLE2 in mammalian cells is also observed during parasite infection in culture and mice.

**Figure 6.**
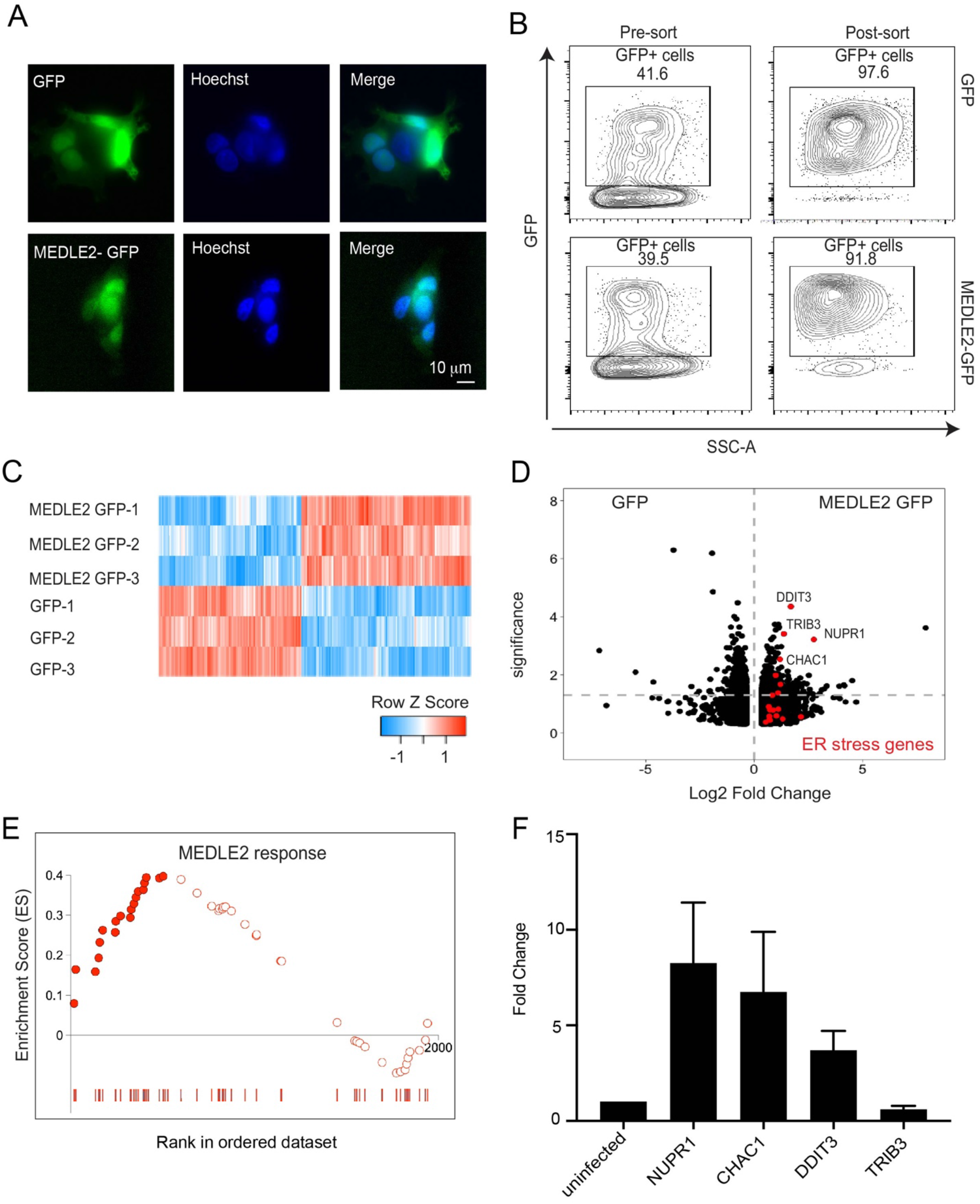
MEDLE2 expressing cells exhibit upregulation of genes involved in the unfolded protein response. (A) HEK293T cells were transfected with plasmids encoding MEDLE2-GFP or GFP alone. After 24h, cells were fixed and processed for IFA. GFP is shown in green, Hoechst in blue. (B) 24h post transection, HEK293T cells were trypsinized and double sorted for live, GFP+ singlets directly into RNA lysis buffer and subjected to RNA sequencing. (C) Heat map depicting the differential gene expression between MEDLE2-GFP (top panel) and GFP control expressing cells (bottom panel). Upregulated gene expression is shown in red (row Z score > 0), while blue shows genes that are downregulated in expression (row Z score < 0). (D) Volcano plot representing the differentially expressed genes found in MEDLE2-GFP expressing cells compared to GFP control cells. 413 transcripts showed upregulation in MEDLE2-GFP expressing cells (right) and 487 genes had lower transcript abundance (left). The horizontal dashed line indicates p-value equal to 0.05. GSEA performed on the 900 differentially expressed genes from the MEDLE2 transfection dataset identifies core enrichment of 20 genes that belong to ER stress response signaling pathways, which are indicated on the volcano plot in red. The most upregulated genes are identified by their gene ID. (E) The 234 genes with the greatest differential expression (*p* <0.01, log fold change absolute value > 1.5) were used to define a MEDLE2 gene set from the MEDLE2-GFP transfection dataset. This signature was used to perform gene set enrichment analysis using data from single cell RNA sequencing on *C.parvum* infected organoid derived cultures, which showed enrichment of 51 genes with 22 genes in the core enrichment for the MEDLE2 response set highlighted in solid red. We note that we did not detect the MEDLE2 response signature in datasets from other enteric infections including Rotavirus (Figure 6-figure supplement 1). (F) Ileal sections were removed from *C. parvum* infected *Ifng^-/-^* mice and uninfected controls (each *n* = 3) and expression levels for the four differentially upregulated genes in the MEDLE2 response set (NUPR1, CHAC1, DDIT3 and TRIB3) were measured by qPCR.

## DISCUSSION

Intestinal cryptosporidiosis in animals and humans is caused by parasites that are morphologically indistinguishable, and therefore were initially described as a single taxon, *C. parvum*. Extensive population genetic studies have since replaced *C. parvum* with a multitude of species, sub species, and strains (Checkley et al., 2015; Xiao et al., 1999). The genomes of these parasites reflect the overall high degree of similarity in their conservation of gene content and synteny; nonetheless, these parasites show pronounced differences in their host specificity (Feng et al., 2017; Feng et al., 2018; Nader et al., 2019). The genomic differences observed are focused on families of predicted secretory proteins that have been proposed to contribute to host specificity, prominently among them the MEDLE family (Fei et al., 2018; Li et al., 2017; Su et al., 2019). Previous studies found that treatment of sporozoites with antisera raised against recombinant MEDLE1 or MEDLE2 diminished the efficiency of infection by 40%. This led the authors to consider that MEDLE proteins may play a role in invasion and that host specificity is rooted in receptor specificity (Fei et al., 2018; Li et al., 2017). In this study, we screened polymorphic genes for the localization of the proteins they encode by epitope tagging the endogenous loci. We found that *C. parvum* exports multiple members of the MEDLE protein family into the cytoplasm of the host cell. This is particularly robust for the highly expressed MEDLE2 protein, where the observation is based on more than 15 independent transgenic strains using different epitope tags and antibodies, in locus and ectopic tagging, and held true in cultured cells and infected animals. Importantly, mutation of the tagged gene changed the localization and molecular weight of the protein, highlighting the specificity of the reagents used.

Apicomplexa evolved multiple mechanisms to deliver proteins to their host cells during and following invasion. The timing of MEDLE2 expression and export and its sensitivity to BFA treatment argues for a mechanism that becomes active after the parasite has established its intracellular niche, in contrast to injection during invasion from organelles already poised to secrete. In *P. falciparum* such export relies on the translocon complex PTEX (de Koning-Ward et al., 2009), which uses the ATPase activity of a heat shock protein to unfold and then extrude cargo through a transmembrane channel into the red blood cell (Ho et al., 2018; Matthews et al., 2019). Genetic ablation of this complex results in loss of export and loss of parasite viability (Beck et al., 2014; Elsworth et al., 2014). In *T. gondii* proteins are exported during intracellular growth via the MYR complex, which is independent of ATP hydrolysis and requires intrinsically disordered cargo (Hakimi et al., 2017). Translocon and export are dispensable for growth of this parasite in culture, but loss profoundly impacts virulence *in vivo* (Hakimi et al., 2017; Marino et al., 2018). With the exception of a putative HSP101 chaperone, genome searches do not readily identify *C. parvum* homologs of the components that make up these two previously characterized translocons (Pellé et al., 2015), but nonetheless, there are important parallels that suggest conserved features of Apicomplexan export.

We demonstrate that MEDLE2 export depends on an N-terminal signal peptide and a leucine residue at position 35. Point mutation of this residue ablates export and results in a higher apparent molecular weight of the protein, which we interpret as a lack of processing at the mutated site. Disorder may be critical for MEDLE2 export, as fusion of well folded domains potently blocked its export into the host cell. This is consistent with an export mechanism unable to unfold proteins as proposed for *T. gondii* and *Plasmodium* liver stages (Beck and Ho, 2021; Gehde et al., 2009; Marino et al., 2018).

Export of proteins in *P. falciparum* and *T. gondii* similarly requires a host targeting motif (Coffey et al., 2015; Hiller et al., 2004; Marti et al., 2004). While the sequence of the motifs varies between species, all share a required leucine residue that has been linked to processing by an aspartyl protease to license export (Boddey et al., 2010; Coffey et al., 2015; Hammoudi et al., 2015; Russo et al., 2010). This protease localizes to the ER in *P. falciparum* where it appears to replace the activity of signal peptidase for exported proteins, while the *T. gondii* homolog acts in the Golgi (Coffey et al., 2015; Marapana et al., 2018). Our mutational analysis in *C. parvum* suggests that cleavage of L35 could replace the activity of signal peptidase similar to *P. falciparum*. The *C. parvum* genome encodes six putative aspartyl proteases, three of which share significant similarity with the Plasmepsin V/ASP5 enzymes responsible for processing exported proteins in *P. falciparum* and *T. gondii*. Ablating these genes may provide an opportunity to block export to the host cell.

Pathogens inject a wide array of factors into the cells of their hosts to remodel cellular architecture, hijack death and survival pathways, change cell physiology and metabolism, rewire transcriptional networks. Many of them aid immune evasion and the evolutionary arms race between host and pathogen and are potent modulators of host specificity (Long et al., 2019; van der Does and Rep, 2007). Here we have identified the first examples of host-targeted effector proteins for intracellular stages of *C. parvum* and investigated the functional role of MEDLE2 during infection. We introduced the protein into its target compartment, the mammalian cytoplasm, by transient transfection and mRNA sequencing revealed a transcriptional signature of ER stress and unfolded protein response (UPR) upon expression of MEDLE2, but not in matched control transfections. This signature is also detectable when cells or mice are infected with *C. parvum*.

ER stress is a common feature of intracellular infection that greatly modulates host cell survival and inflammatory response. Bacteria, viruses and protozoa alike have evolved strategies to trigger or block the UPR during infection suggesting the response not to be a byproduct of infection, but rather an active participant (Abhishek et al., 2018; Alshareef et al., 2021; Perera et al., 2017). This relationship is complex and depending on cellular context, the UPR can aid pathogens or restrict them. Modulating the UPR is pivotal for intracellular bacteria that reside in vacuoles, including *Chlamydia, Brucella*, *Listeria,* and *Mycobacterium,* to build their intracellular niche and to acquire nutrients (Celli and Tsolis, 2015). *Cryptosporidium* lost the ability to synthesize most required metabolites and depends on an elaborate membranous feeder organelle at the host-parasite interface. This reduction includes biosynthesis of many lipids (Coppens, 2013) and we note modulation of lipid synthesis and mobilization as one important consequence of the UPR (Moncan et al., 2021). The UPR is also important in the context of recognition of intracellular pathogens and the subsequent cytokine-mediated immune responses. *Cryptosporidium* triggers an enterocyte intrinsic inflammasome leading to the release of IL-18 from the infected cell (McNair et al., 2018) and infection also results in pronounced production of interferon λ (Ferguson et al., 2019). It remains to be determined whether the UPR is a byproduct of host detection of the disordered protein MEDLE2 following its translocation during *C. parvum* infection, or if it is an active process deliberately triggered to promote parasite survival.

The MEDLE genes are highly polymorphic and encoded by multiple loci, but our current view of the family likely underestimates its true plasticity. The recent reannotation of the *C. parvum* genome using long read sequencing found evidence for multiple ‘alternative’ telomeres carrying different MEDLE gene clusters (Baptista et al., 2021). Our experimental observations support this, as when we ablated the MEDLE2 gene by homologous recombination, additional MEDLE2 copies remained in the genome (Figure 1-figure supplement 2). Similar observations were made when tagging the gene (Fig1B, Figure2-figure supplement 1). It is important to note that *Cryptosporidium* has a single host lifecycle and undergoes sexual exchange throughout the infection, providing near constant opportunity for recombination (Tandel et al., 2019). This is reminiscent of the importance of chromosomal position and recombination for the antigenically varied VSG and var genes in *Trypanosoma brucei* and *P. falciparum*, respectively (Dreesen et al., 2007; Zhang et al., 2019). The fact that MEDLE proteins are delivered into the host cell makes them prime targets for immunity, and immune evasion may drive chromosomal organization, expression and evolution of these genes.

## ACKNOWLEDGEMENTS

This work was supported in part by funding from the National Institutes of Health through grants to BS (R01AI127798 and R01AI112427), and fellowships and career awards to JED (T32AI007532), AS (K99AI137442), and JAG (T32A1055400). We thank Andrea Stout, Jasmine Zhao (Cell and Developmental Biology Microscopy Core), Gordon Ruthel (Penn Vet Imaging Core), Dan Beiting (Center for Host Microbial Interactions), and Patty Costello (Comparative Pathology Core) for help with microscopy, RNA sequencing, and histology, respectively. We thank Jacek Gaertig and Phillip Scott for sharing reagents.

## AUTHOR CONTRIBUTIONS

JED developed protein export assays, characterized MEDLE2 expression and assessed the requirements for MEDLE2 export through mutant studies with support from AGR, AS and EH. AS helped prepare intestinal samples for IHC. JAG and JTC assisted with flow cytometry. ARG performed the bioinformatics analysis. JED, AS, and BS conceived of the study and JED and BS wrote the manuscript.

## MATERIALS AND METHODS

### Contact for reagent and resource sharing

For access to reagents or parasite strains used in this study please contact Dr. Boris Striepen: Tel.: 1-215-573-9167; Fax: 1-215-746-2295.

### Mouse models of infection

*Ifng^-/-^* (stock no:002287) were purchased from Jackson Laboratory and maintained as a breeding colony at the University of Pennsylvania. All mice used in this study ranged from 4 – 12 weeks of age. We note that both male and female *Ifng^-/-^* mice were used to generate and propagate *C. parvum* transgenic parasite lines, without a difference being noted in parasite shedding. All protocols for animal care were approved by the Institutional Animal Care and Use Committee of the University of Georgia (protocol A2016 01-028-Y1-A4) and the Institutional Animal Care and Use Committee of the University of Pennsylvania (protocol #806292).

### Parasite strains

*Cryptosporidium parvum* transgenic strains were made and propagated in *Ifng^-/-^* mice (stock no:002287). Oocysts were then purified from fecal collections using sucrose flotation followed by a cesium chloride gradient (see Method details). All *Cryptosporidium parvum* oocysts used in this study as WT controls, as well as to generate transgenic strains, are on the IOWAII strain background, purchased from Bunchgrass Farms (Dreary, ID).

### Plasmid construction

Guide oligonucleotides (Sigma-Aldrich, St. Louis. MO) were introduced into the *C. parvum* Cas9/U6 plasmid by restriction cloning, as detailed in (Pawlowic et al., 2017). All plasmids encoding epitope tags, as well as for ectopic MEDLE2 expression were constructed by Gibson assembly using NEB Gibson Assembly Master Mix (New England Biolabs, Ipswich. MA). A linear repair template was generated by PCR. See Supplementary Table 2 for a complete list of primers used for this study.

### Generation of transgenic parasites

Transgenic parasites were derived as previously described (Sateriale et al., 2020). Briefly, 5 × 10^7^ *C. parvum* oocysts were bleached on ice, washed in 1X PBS and incubated in sodium taurodeoxycholate. Excysted sporozoites were resuspended in transfection buffer supplemented with a total of 100 µg DNA (comprised of 50 µg of Cas9/gRNA plasmid and 50 µg of repair template generated by PCR) and nucleofected using an Amaxa 4D nucleofector (Lonza, Basel, Switzerland). Transfected parasites were resuspended in PBS and administered to *Ifng^-/-^* mice. Mice were pretreated with antibiotics for 1 week preceding infection and with sodium bicarbonate immediately before parasite administration (Sateriale et al., 2020). Mice received 16mg/mL Paromomycin in drinking water for selection. Transgenic parasites were detected by measuring fecal nanoluciferase activity and purified from feces using sucrose flotation followed by a cesium chloride gradient and stored in PBS at 4°C (Sateriale et al., 2020).

### Nanoluciferase Assay to monitor parasite shedding

20 mg of fecal material was dissolved in nanoluciferase lysis buffer and mixed 1:1 with nanoluciferase substrate/nanoluciferase Assay Buffer (1:50) in a white bottom plate. Relative luminescence was read using a Promega GloMax Plate Reader.

### Integration PCR to confirm generation of transgenic parasites

DNA was purified from excysted sporozoites using the Qiagen DNeasy Blood and Tissue kit (Qiagen 69504). PCR primers were designed to anneal outside of the 5’ and 3’ homology arms used to direct homologous recombination and matched with primers annealing to the nanoluciferase reporter gene or the Neomycin selection marker respectively. Primers for the thymidine kinase gene served as control, unless otherwise noted. Where indicated, amplicons were cloned using the ZeroBlunt TopoTA kit (Invitrogen 450245) and transformed into One Shot^TM^ Topo10 Chemically Competent *E. coli* (Invitrogen C404003). Individual colonies were miniprepped and sequenced.

### *In vitro* infection and Immunofluorescence Assay

Coverslips seeded with human ileocecal adenocarcinoma cells (HCT8) (ATCC® CCL-244™) were infected when 80% confluent with 200,000 purified oocysts (bleached, washed, and resuspended in RPMI medium containing 1% serum). For time course infections, parasites were allowed to invade for 3 hours, then medium was removed, and the cells were washed with PBS to remove unexcysted oocysts and replaced with fresh RPMI medium with 1% serum. At indicated time points, cells were washed with PBS, and successively fixed and permeabilized with PBS supplemented with 4% paraformaldehyde or 0.1% Triton X-100 for 10 min each (Sigma St. Louis, MO). Coverslips were blocked with 1% Bovine Serum Albumin (BSA) (Sigma St. Louis, MO). Antibodies were diluted in blocking solution. The was rat monoclonal anti-HA (Millpore Sigma, Burlington, MA, USA) was used as primary antibody (1:500) and goat-anti-rat polyclonal Alexa Fluor 594 (Thermofisher, Waltham, MA) as secondary along with *Vicia villosa* lectin (Vector Labs Burlingame, CA, USA). Host and parasite nuclei were stained with Hoechst 33342 (Thermofisher, Waltham, MA). Slides were imaged using a Zeiss LSM710 Confocal microscope or a Leica Widefield microscope. Experimental slides were prepared and imaged in duplicate for a minimum of 2 biological replicates.

### Immunohistochemistry on infected intestine

Infected *Ifng^-/-^* mice were euthanized at day 12 during peak infection, and the distal 1/3 of the small intestine was dissected. The tissue was washed with PBS and ‘swiss-rolled’ and fixed overnight in 4% Paraformaldehyde at 4°C, placed in 30% sucrose in PBS for cryoprotection, and mounted with OCT compound (Tissue-Tek, Sakura Finetek, Japan) and frozen. Cryomicrotome sections were permeabilized and blocked and labeled as described above (Alexa Fluor 647 Phalloidin (Thermofisher, Waltham, MA) was used in addition). Sections were prepared in duplicate and imaged using a Zeiss LSM710 Confocal Microscope.

### Poly-L-Lysine treatment of coverslips and sporozoite IFA

Sterile coverslips were treated with Poly-L-Lysine (Sigma, St Louis, MO, USA), washed with water for 5 min and airdried. Sporozoites suspended in PBS were allowed to settle on treated coverslip for 1h prior to fixing and IFA. Primary antibodies used were mouse anti-Cp23 (1:100) (LS Bio Seattle, WA) and rat monoclonal anti-HA (1:500). Secondary antibodies include: goat-anti-rat polyclonal Alexa Fluor and goat-anti-mouse polyclonal Alexa Fluor 488 (both 1:1000, Thermofisher, Waltham, MA).

### Brefeldin A (BFA) treatment during *C. parvum* infection

Excysted oocysts were allowed to invade HCT-8 coverslip cultures for 3h, unexcysted oocysts were removed by PBS wash, and cultures we replaced with medium supplemented with 1% serum and 10 µg/mL BFA from a 1000X stock in DMSO. Medium supplemented with carrier alone served as control. Cultures were fixed and processed 10h post infection.

### Live imaging of Beta-Lactamase Reporter Assay

1 ×10^6^ WT and MEDLE2-BLA oocysts were used to infect HCT8 cells in a 35 mm glass bottom dish (MatTek Life Sciences Ashland, MA). After 24h, the medium was replaced with RPMI medium containing CCF4-AM substrate from the LiveBLAzer^TM^ FRET-B/G Loading Kit (ThermoFisher Scientific Waltham, MA). Cells were incubated in the dark at 37°C for 1h, washed with PBS 3 times and live imaged using a Leica SP5 Confocal Microscope using a water immersion lens.

### Cre recombinase reporter assay by flow cytometry

Pre-made lentivirus was used to transform HCT8 cells with a loxP GFP/RGP color switch cassette (GenTarget Inc San Diego, CA). Cells were selected with 400 mg/mL Neomycin (Millpore Sigma, Burlington, MA) for 14 days and validated by transfection with 5 µg Cre recombinase plasmid using LipofectamineP3000 (ThermoFisher Scinetific Waltham, MA). After 24h and 48h, cells were trypsinized and flow sorted using a LSRFortessa (BD Biosciences) and data were analyzed with FlowJo v10 software (TreeStar). 1 × 10^6^ WT and MEDLE2-Cre oocysts were used to infect 6 well cultures of Lox GFP/RFP color switch cells. After 48h, cells were trypsinized and resuspended in 1 mL PBS. 300 µL were used for Nanoluciferase assay and 700 µL cells for flow cytometry. Forward and side scatter was used to gate viability, untransfected uninfected cells to establish the green gate, and Cre recombinase transfected cells for the red gate (3 biological replicates for each condition).

### Western Blot on *C. parvum* Infected Cells

HCT8 cultures infected with 5 × 10^6^ oocysts were for 48h were treated with Trypsin-0.25% EDTA (ThermoFisher Scientific, Waltham, MA), pelleted, and flash frozen in liquid nitrogen. Cell pellets were lysed in Pierce^TM^ IP Lysis Buffer (ThermoFisher Scientific, Waltham, MA), supplemented 1:100 with both protease inhibitor cocktail (Sigma St. Louis, MO) and benzonase nuclease (Millpore Sigma Burlington, MA). Lysates were incubated on ice for 15 min, sonicated (80% amplitude, 10 second pulses, rest on ice for 1 min between 3 times) cleared by centrifugation (20,000 g, 10 min, 4°C), mixed with freshly prepared Lamelli Sample buffer (Millpore Sigma Burlington, MA) + β-Mercaptoethanaol (1:20) (Sigma St. Louis, MO), boiled and loaded on a 12% Mini-PROTEAN® TGX™ Precast Protein Gel (BioRad Hercules, CA) run at 70 V for 2.5h. Gels were transferred to 0.45 µm pore size Nitrocellulose membrane (ThermoFisher Scientific Waltham, MA) overnight at 0.02 A at 4°C. The membrane was blocked for 1h with Intercept (TBS) Protein-Free Blocking Buffer (LI-COR Lincoln, NE), antibodies we diluted in blocking solution with 0.01% Tween®20 (Sigma, St. Louis, MO) using rat monoclonal anti-HA 1:500 (Millpore Sigma, Burlington, MA) and rabbit anti-neomycin phosphotransferase II 1:1000 (Millpore Sigma, Burlington, MA) as primary and IRDye® 800CW Goat anti-Rat IgG and IRDye® 680RD Goat anti-Rabbit IgG (both 1:10,000, LI-COR, Lincoln, NE) as secondary antibody. Washed membranes were imaged using a Odyssey Infrared Imaging System v3.0 (LICOR, Lincoln, NE).

### Generation of MEDLE2 mutant plasmids for host cell transfection

Human codon optimized MEDLE2 lacking the N terminal signal peptide (aa 21-209) was synthesized by Integrated DNA Technologies (IDT, Coralville, IA) and cloned into the mEGFP-Lifeact-7 mammalian expression plasmid (Addgene #54610), replacing Lifeact and appending a 3x HA tag. Point mutations were engineered by Gibson cloning. HEK293T cells (ATCC®CRL-3216^TM^) were transfected with 5 µg of each plasmid using Lipofectamine®P3000 (ThermoFisher Scientific Waltham, MA). 24h post transfection, cells were harvested and processed for Western Blot analysis.

### Flow Cytometry analysis of transfected cells

HEK293T cells were subjected to lipofection with 25 µg GFP only plasmid or MEDLE2-GFP plasmid, grown for 24h, trypsinized, washed and resuspended in PBS with DAPI and passed through a 40 µM filter (BD Biosciences San Jose, CA). Cell viability was gated based upon DAPI staining. Untransfected HEK293T served as negative control and GFP expressing HEK293T cells as positive control to establish gates. 10,000 green, single cells were double sorted using an Aria C flow cytometer first into PBS then into lysis buffer (3 biological replicates for each condition).

### RNA extraction sequencing and data analysis

Total RNA was extracted using the Qiagen RNeasy Microkit (Qiagen, Germantown, MD) and input RNA was quality controled and quantified using a Tape Station 4200 (Aglient Technologies Santa Clara, CA). cDNA synthesis was performed following the clonTechSMART-seq cDNA synthesis protocol (15 cycles). Following DNA cleanup, a Nextera library was prepared and nucleic acid was quantified using the Qubit 3 Fluorometer (Thermo Fisher Scientific Waltham, MA). Samples were pooled for RNAsequencing of 4 nM of total cDNA and sequencing was performed using a NextSeq 500 Instrument (Illumina Inc San Diego, CA).

RNAseq reads were pseudo-aligned to the Ensembl *Homo sapiens* reference transcriptome v86 using kallisto v0.44.0 (Bray et al., 2016). In R, transcripts were collapsed to genes using Bioconductor tximport (Robinson et al., 2010) and differentially expressed genes were identified using Limma-Voom (Law et al., 2014; Ritchie et al., 2015). The MEDLE2 transcription response data set can be found under GEO accession number GSE174117. Gene set enrichment analysis (GSEA) was performed using the GSEA software and the annotated gene sets of the Molecular Signatures Database (MSigDB) (Mootha et al., 2003; Subramanian et al., 2005). The MEDLE2 signature was generated from the differentially expressed genes and read into GSEA to evaluate its presence in published datasets of *C. parvum infection* (Nikolaev et al., 2020; Saxena et al., 2017).

### qPCR for MEDLE2 response genes

8-week-old *Ifng^-/-^* mice were infected with 10,000 *C. parvum* oocysts and the infection was tracked by fecal nanoluciferase activity. Infected mice (*n* = 3)and uninfected controls (*n* = 3) were euthanized after 10 days and the distal 1/3 of the small intestine was removed. The tissue was washed with 1X PBS until clear of fecal material and then cut longitudinally. 5 mm diameter gut punches were made and preserved in RNA*late*r^TM^ solution (Thermo Fisher Scientific Waltham, MA). RNA was extracted from tissue samples using the Qiagen RNeasy MiniKit (Qiagen, Germantown, MD) following homogenization with a bead beater and passage through a QIAshredder (Qiagen, Germantown, MD). 5 µg of cDNA was reverse transcribed using the SuperScript^TM^ First Strand Synthesis kit following the manufacturer’s instructions for use with OligoDT (Thermo Fisher Scientific Waltham, MA). qPCR was performed using a Viia^TM^7 Real-time PCR System (Thermo Fisher Scientific Waltham, MA) and relative gene expression was determined using the ΔΔCT method.

### Quantification and Statistical Methods

GraphPad PRISM was used for all statistical analyses. When measuring the difference between two populations, a standard *t*-test was used. For data sets with 3 or more experimental groups, a one-way ANOVA with Dunnett’s multiple comparison’s test was used. Simple Linear Regression was used to determine the goodness of fit curve for the number of MEDLE2 expressing cells and intracellular parasites. Quantification of imaging experiments was performed using ImageJ macros programmed to count both parasites and host cell nuclei in blinded images that were captured using a scanning function to avoid bias during acquisition.

## SUPPLEMENTARY MATERIALS

**Figure 1 Supplemental Table 1:** Members of multigene families for which localization of protein product was initially attempted in this study

| Gene family | Gene ID | Result |
| --- | --- | --- |
| MEDLE | cgd5_4590 | Exported to host cell |
| FLGN | cgd4_4470 | Transgenic unsuccessful |
| GGC | cgd5_3570 | Transgenic unsuccessful |
| SKSR | cgd8_30 | Not in host cell |
| SKSR | cgd8_40 | Not in host cell |
| WYLE | cgd8_3560 | Not in host cell |

**Figure 1- figure supplement 1.**
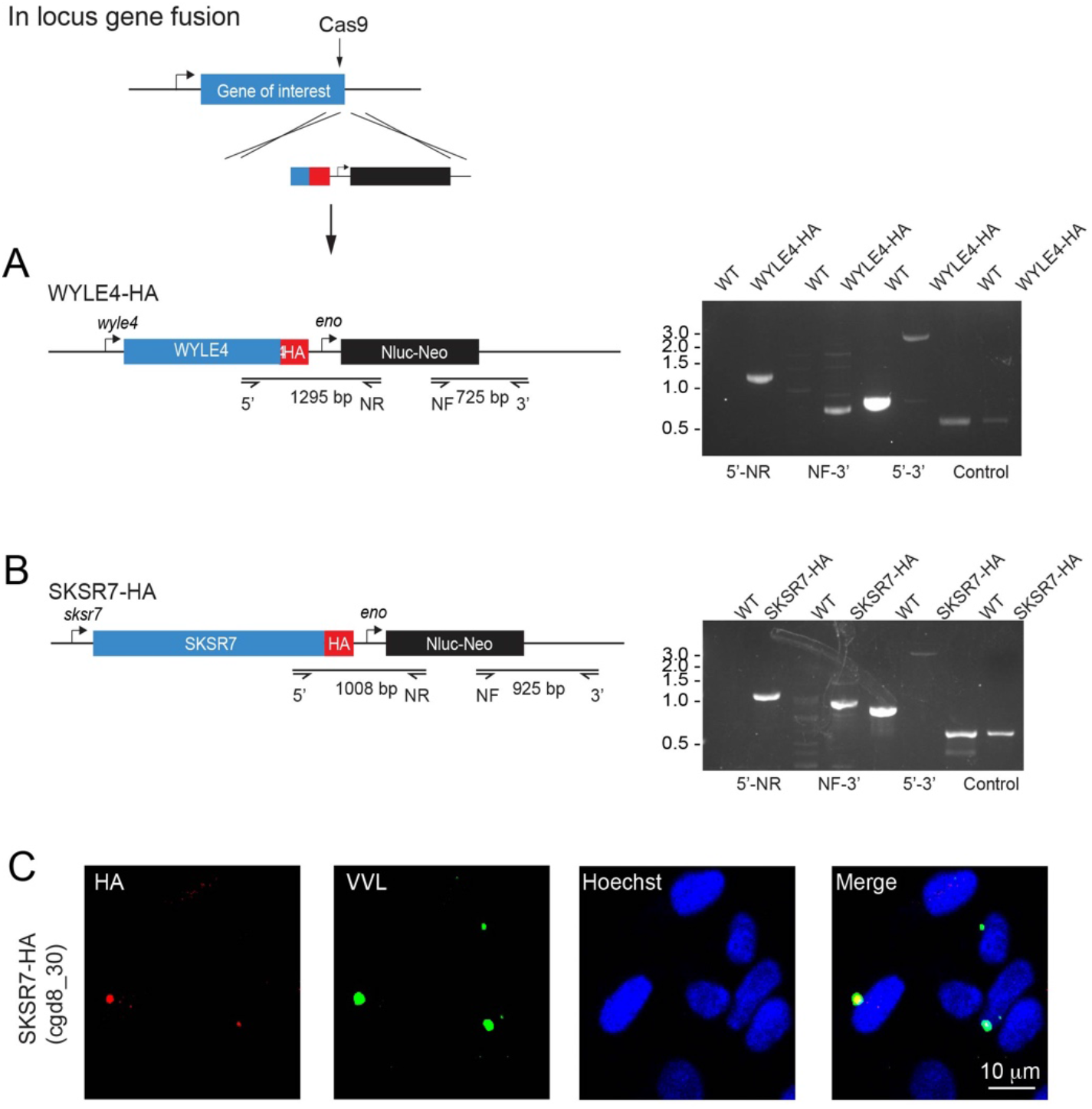
Additional secretory proteins tested in this study. (A-C) Schematic depicting the generation of in locus gene fusion transgenic parasite strains for WYLE4 (A), and SKSR7 (B). In each case, the C terminus of the gene was targeted for integration of a repair construct containing an HA epitope tag, a Nanoluciferase reporter gene (Nluc) and a Neomycin phosphotransferase (Neo) selectable marker. Individual primer pairs used to map integration in WT and transgenic strains are shown. (C) SKSR7-HA does not localize to the host cell; rather, the protein (red) exhibits expression in the parasite (green).

**Figure 1- figure supplement 2.**
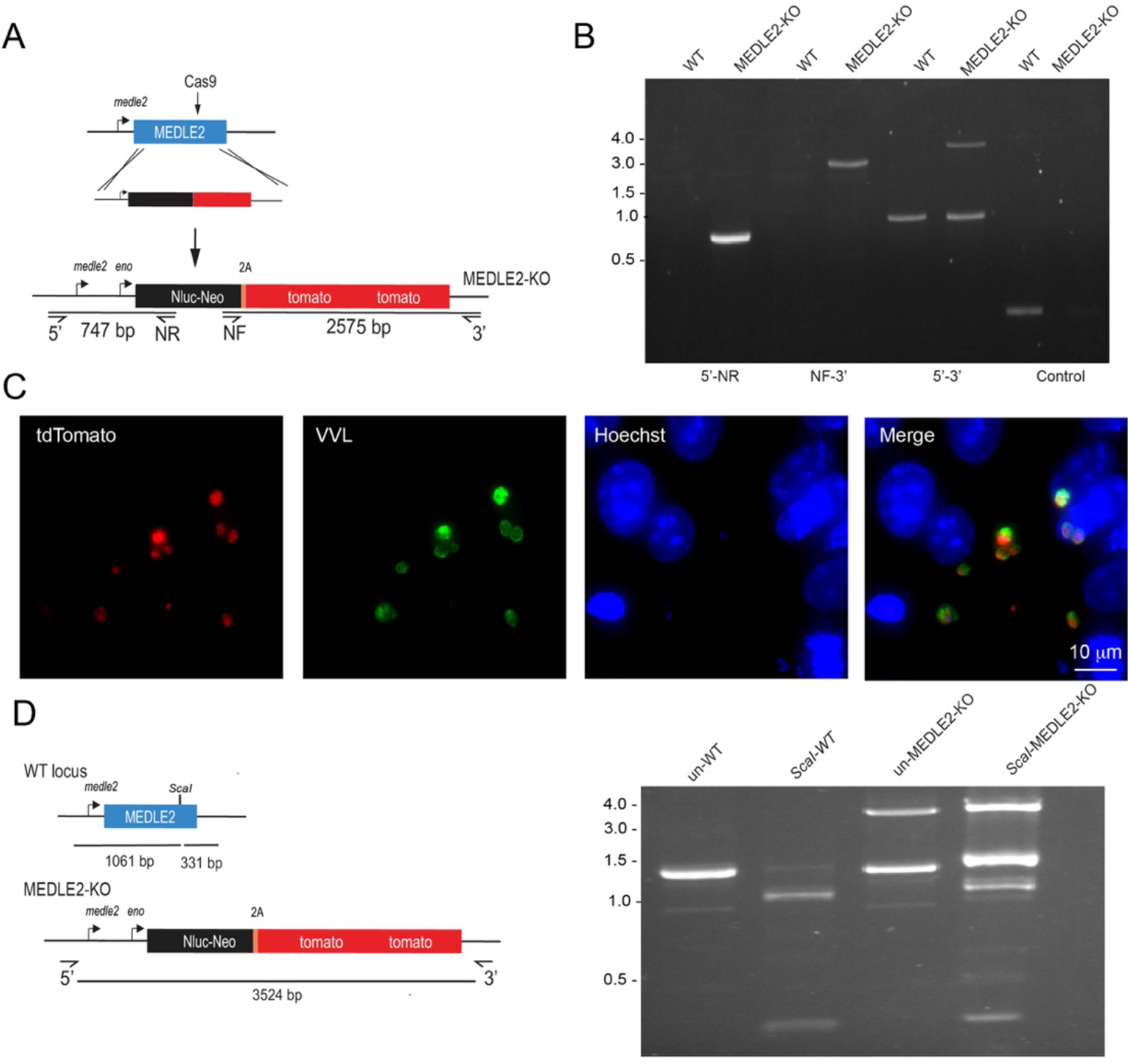
Knockout of MEDLE2 reveals multiple copies of the gene in the genome. (A) Schematic representation for the strategy used to generate a MEDLE2 KO line, in which the entire locus of MEDLE2 is replaced with a nanoluciferase reporter gene (Nluc) and the neomycin phosphotransferase (Neo) selection marker fused to a 2A peptide, and tdTomato, such that the parasites express a red fluorescent protein in their cytoplasm. The solid black arrow indicates the position of the Cas9 induced double-stranded break in the middle of the gene. Note, this is a different guide from the one used for C terminal tagging. (B) PCR mapping modification of the MEDLE2 locus using genomic DNA from wild type (WT) and transgenic (MEDLE2 KO) sporozoites using the primer pairs shown in A and the Thymidine kinase (TK) gene as a control. Note the persistence of a WT band (1392 bp) in the 5’ – 3’ amplification product, despite the presence of the transgene (3524 bp). (C) HCT-8 cultures were fixed 24h after being infected with MEDLE2 KO transgenic parasites. MEDLE2 KO parasites exhibit red fluorescence in their cytoplasm as expected (Red, tdTomato, parasite cytoplasm; green, parasites VVL; blue, Hoechst). (D) The full gene PCR products from WT (1392 bp) and MEDLE2 KO parasites (3524 bp) were used for restriction digest with ScaI. A single ScaI restriction site is found in the C terminus of WT MEDLE2; however, integration of the repair cassette disrupts this site. ScaI digested WT PCR product results in 2 digest products: 331 bp and 1061 bp. Undigested MEDLE2 KO full gene product has the expected 3524 bp fragment, as well as a persisting 1392 bp WT band. ScaI digested MEDLE2 KO, shows the 3525 bp repair cassette resistant to ScaI digest, as well as the 331 bp and 1061 bp fragments produced from digest of the unmodified MEDLE2 locus. As a result, there are multiple copies of MEDLE2 in the genome and we have only targeted one for knockout.

**Figure1- figure supplement 3.**
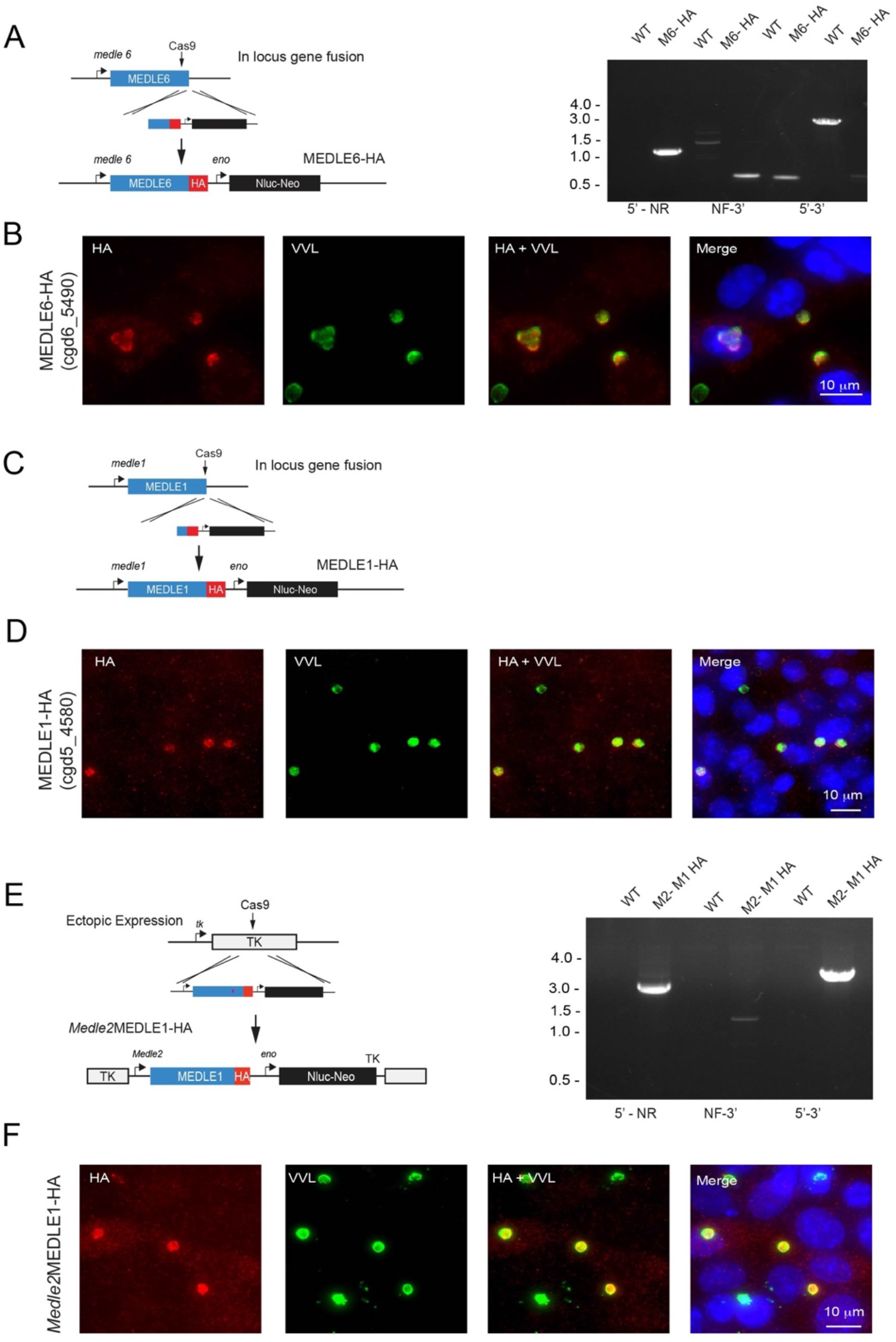
Other members of the MEDLE gene family are exported to the host cell. (A) Schematic representation depicting the generation of a MEDLE6-HA transgenic parasite line, in which the endogenous locus of MEDLE6 (cgd6_5490) is HA epitope tagged at the C terminus. Proper integration at the desired locus was confirmed using PCR mapping with gDNA isolated from MEDLE6 transgenic parasites (M6) and a wild type control (WT). (B) MEDLE6-HA parasites were used to infect HCT8 cells for an immunofluorescence assay. Cells were fixed every 12h and stained for IFA. Shown as a representative image, at 24h post infection, MEDLE6 (red) localizes in/around the parasite (green), as well as slightly in the host cell. Host cell expression is more apparent in multiply infected cells. (C) The MEDLE1 (cgd5_4580) locus was targeted for integration of an HA epitope tag at the C terminus. (D) MEDLE1-HA parasites were used to infect HCT8 cells for a time course infection and IFA was performed on cells fixed every 12h. Shown as a representative image, at 12h post infection, MEDLE1 (red) localizes in/around the parasite (green), as well as at very low levels in the host cell. (E) Schematic representation for the strategy used to engineer a MEDLE1 overexpression line. The MEDLE2 promoter was used to drive expression of an ectopic copy of MEDLE1-HA expressed in the TK locus. Proper integration was assessed using PCR mapping with gDNA isolated from *Medle2*MEDLE1-HA (M2-M1 HA) and WT control (WT) parasites. (F) M2-M1 HA parasites were used to infect HCT8 cells for IFA. At 24h post infection, MEDLE1-HA (red) can be seen in/around the parasite (green), as well as in the host cell when expression is driven by the MEDLE2 promoter.

**Figure 2- figure supplement 1.**
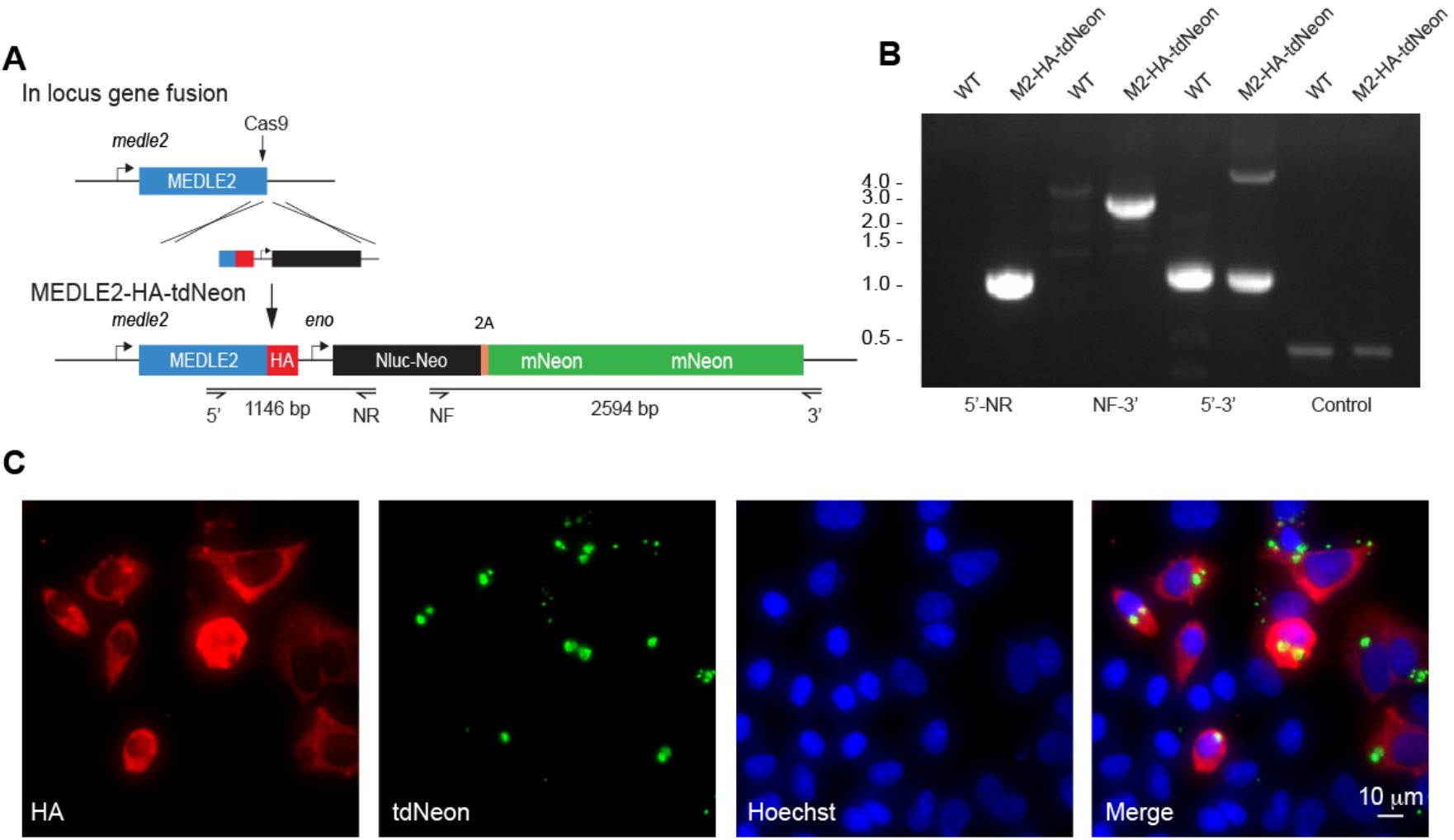
Construction of a MEDLE2-HA cytoplasmic tdNeon reporter parasite. (A) Schematic representation of the strategy to derive reporter parasite line in which MEDLE2 is 3X HA epitope tagged and the parasite cytoplasm expresses a tandem mNeon green tag (tdNeon). The solid black arrow indicates the position of the Cas9 induced double-stranded break at the C terminus of the gene, which is the same guide used in Figure 1B to generate the MEDLE2-HA transgenic parasites. (B) PCR mapping modification of the MEDLE2 locus using genomic DNA from wild type (WT) and transgenic (MEDLE2-HA-tdNeon) sporozoites using the primer pairs shown in A and the Thymidine kinase (TK) gene as a control. Note the presence of both a 1174 bp WT gene and a 4557 bp transgene. (C) HCT-8 cultures were infected with and MEDLE2-HA-tdNeon transgenic parasites and fixed at 24h. Red, HA tagged protein; green, parasites (tdNeon); blue, Hoechst.

**Figure 4- figure supplement 1.**
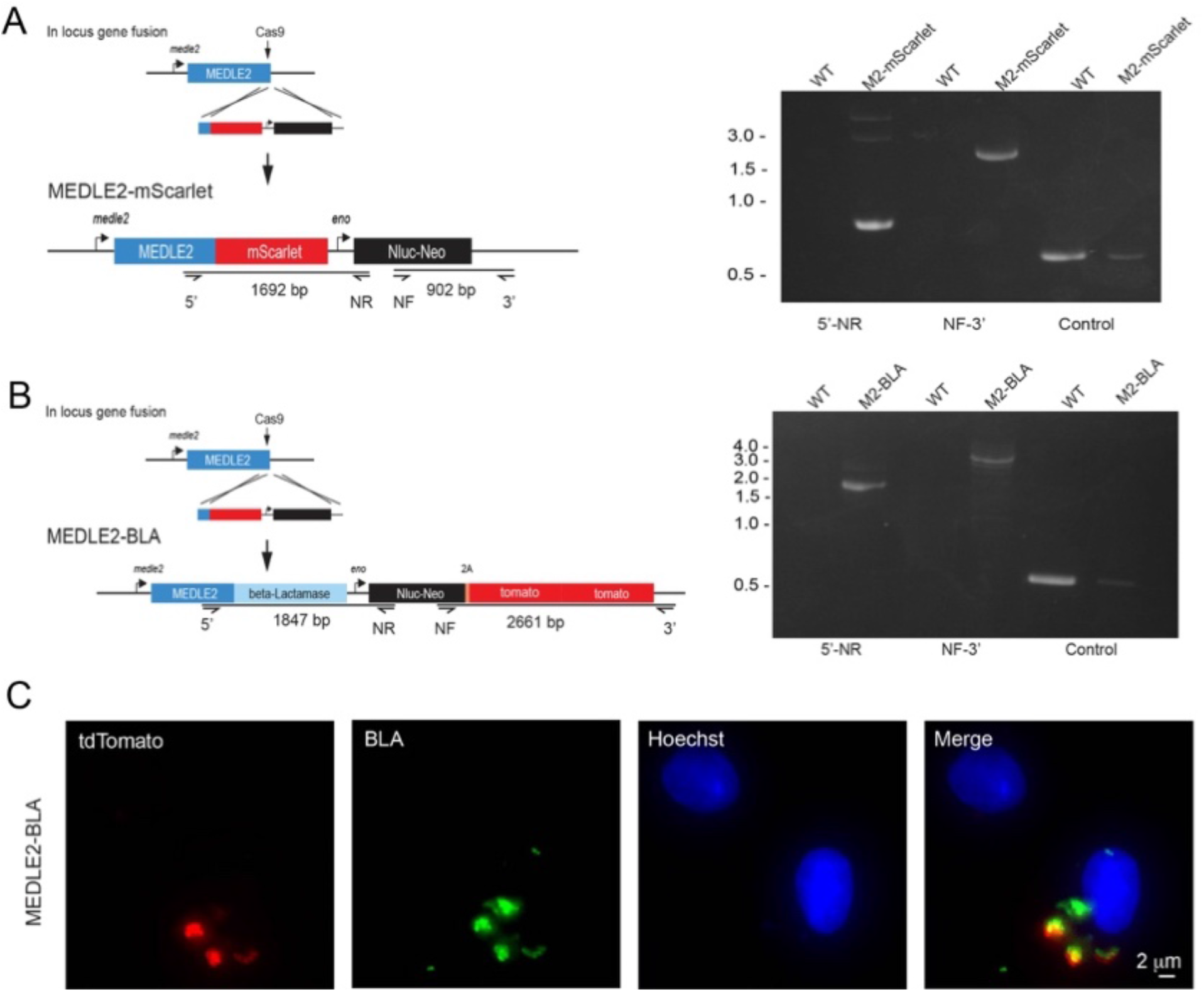
MEDLE2-BLA is not exported. (A) Schematic representation of the strategy used to generate a MEDLE2-mScarlet parasite line, in which the MEDLE2 is C terminally tagged with the fluorescent protein mScarlet, as well as engineered to express a nanoluciferase reporter gene (Nluc) and the neomycin phosphotransferase (Neo) selection marker. The solid black guide hit sequence is the same as used in Figure 1B to generate the MEDLE2-HA transgenic parasites. Integration PCR using MEDLE2-mScarlet and WT gDNA confirms proper integration. (B) The C terminus of MEDLE2 was targeted for insertion of a construct encoding Beta-lactamase (BLA), a nanoluciferase reporter gene (Nluc) and the neomycin phosphotransferase (Neo) selection marker fused to a 2A peptide and a tdTomato reporter gene, using the same tagging strategy previously utilized for this locus. Integration PCR mapping the MEDLE2 locus using genomic DNA from wild type (WT) and transgenic (MEDLE2-BLA) sporozoites with the corresponding primer pairs shown in A and the Thymidine kinase (TK) gene as a control shows the locus was successfully modified. (C) MEDLE2-BLA transgenic parasites were used to infect HCT8 cells and fixed at 24h for IFA. Red, parasites (tdtomato); green, Beta-lactamse (Beta-lactamase antibody); blue, Hoechst.

**Figure 4- figure supplement 2.**
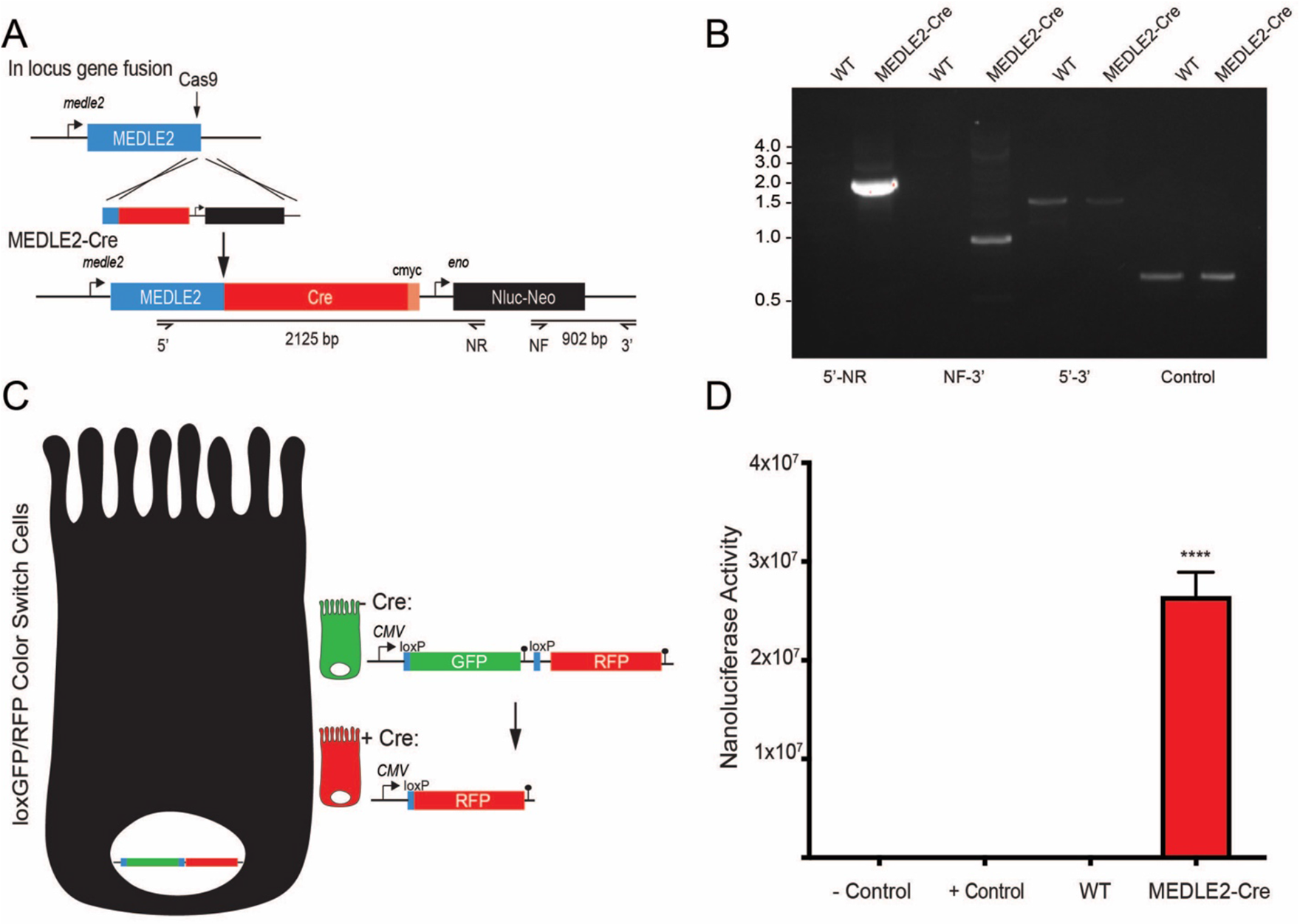
MEDLE2-Cre parasites infect loxGFP/RFP color switch cells. (A) MEDLE2-Cre expressing parasites were generated by targeting the MEDLE2 locus for insertion of a construct encoding Cre recombinase (Cre), a nanoluciferase reporter gene (Nluc) and the neomycin phosphotransferase (Neo) selection marker. The solid black arrow indicates the guide hit sequence, which is the same guide used in Figure 1B to generate the MEDLE2-HA transgenic parasites. (B) Integration PCR mapping the MEDLE2 locus using genomic DNA from wild type (WT) and transgenic (MEDLE2-Cre) sporozoites using the primer pairs shown in A and the Thymidine kinase (TK) gene as a control. (C) Schematic of the LoxGFP/RFP color switch HCT8. Introduction of Cre into these cells induces a color switch from green to red by recombinase directed removal of the GFP coding sequence along with a stop codon preventing translation of RFP. (D) 1 × 10^6^ transgenic MEDLE2-Cre oocyts were used to infect LoxGFP/RFP color switch HCT8 cells for 48h before preparation of the sample for flow cytometry. 400,000 cells were removed from the sample preparation for nanoluciferase assay to determine infection in the culture. Mean nanoluciferase (relative luminescence) ± SEM is shown for three replicates (*p* < 0.0001; one-way ANOVA with Dunnett’s Multiple Comparison test). Uninfected cells were used as a negative control and Cre recombinase transfected cells were used as a positive control.

**Figure 5- figure supplement 1.**
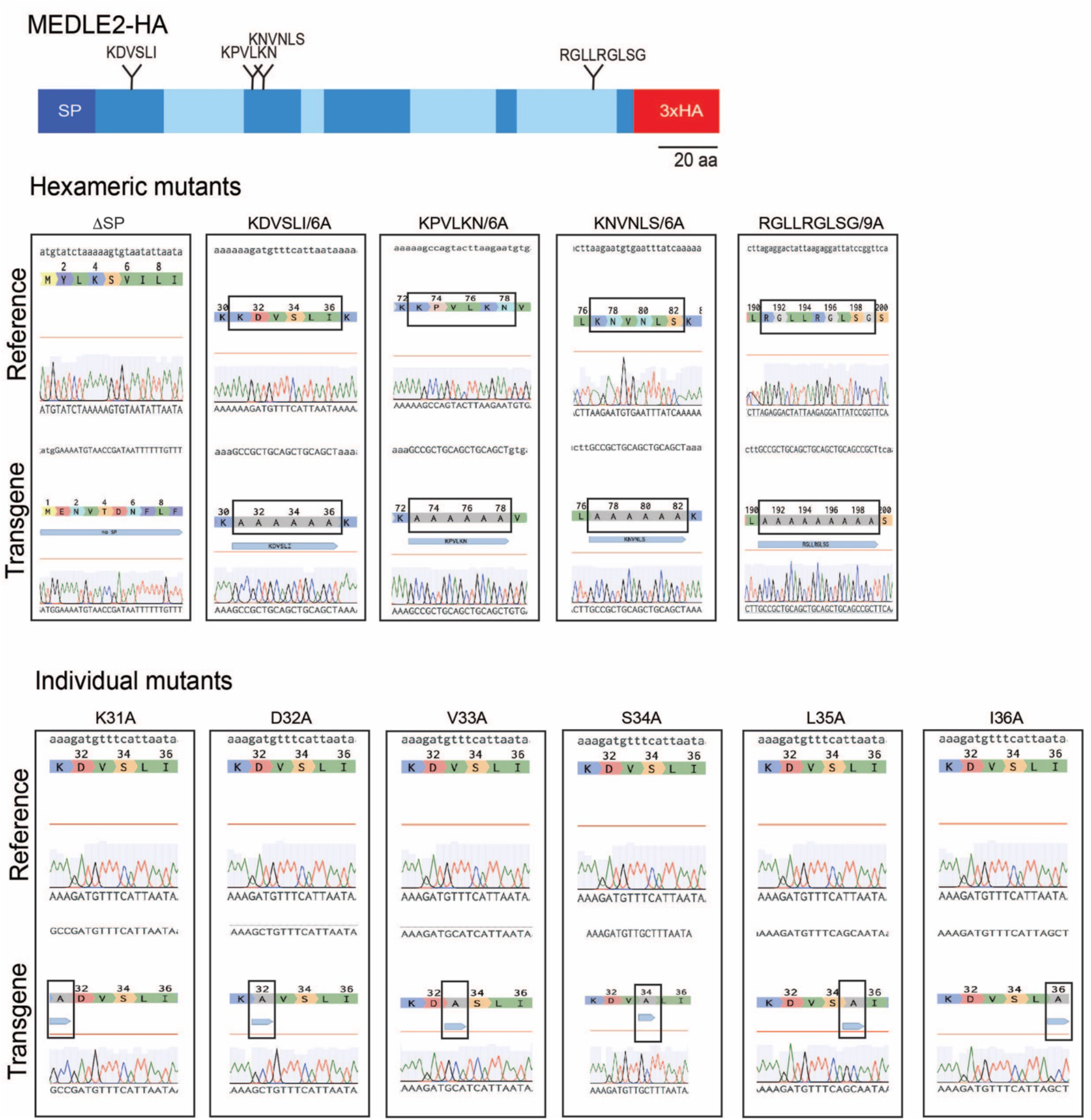
Sanger sequencing confirming the generation of MEDLE2 mutants. 100,000 sporozoites were used from each mutant strain for genomic DNA extraction. The resulting gDNA was used for PCR mapping of the TK locus to verify the desired mutagenesis. 5’ TK – Nluc PCR products were used for TopoTA cloning and the resulting colonies were grown, and Sanger sequenced. 3 colonies were sequenced for each strain using the M13 forward, M13 reverse, and an internal MEDLE2-specific primer to confirm targeted mutagenesis. The Benchling alignment of the Sanger sequencing result (transgene) to the reference sequence is shown. The black box highlights the mutation engineered in each strain.

**Figure 5- figure supplement 2.**
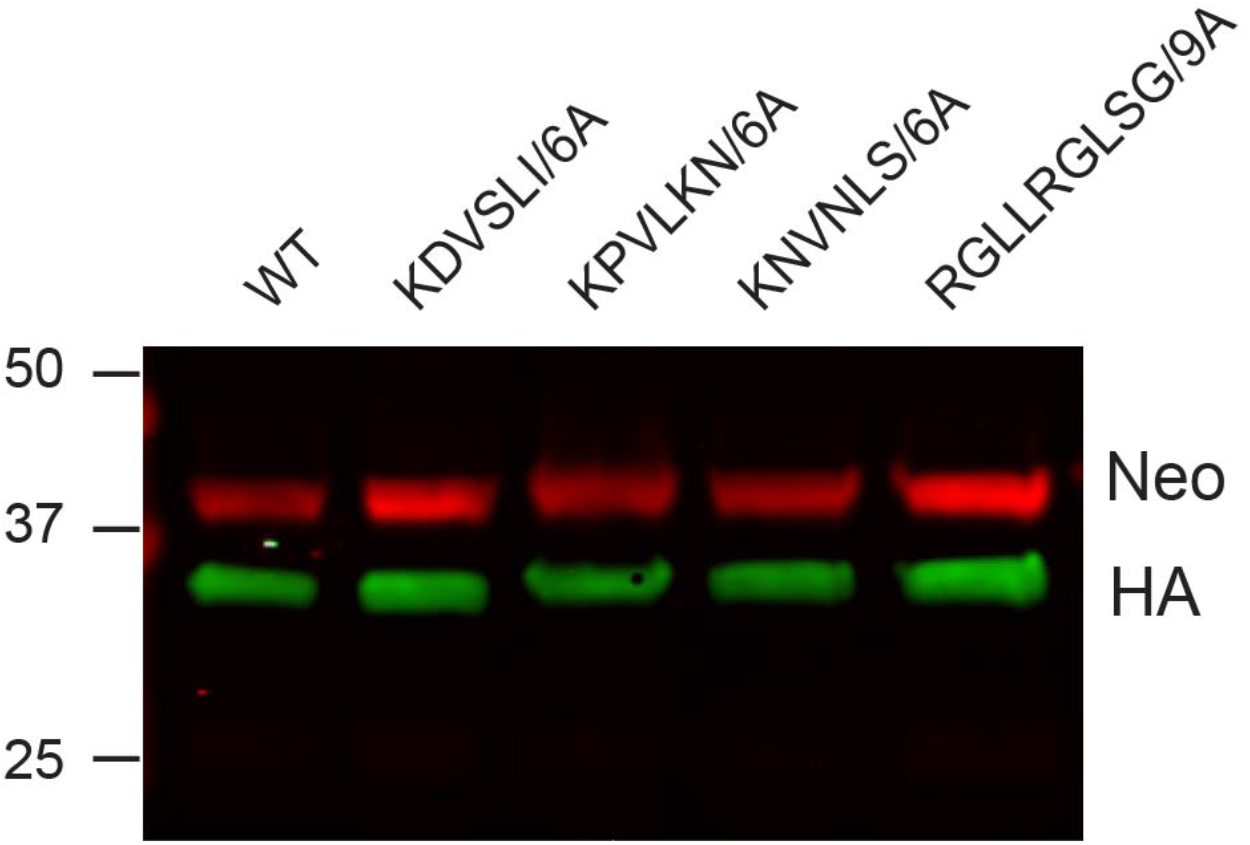
MEDLE2 mutants are of the same size as WT MEDLE2 when expressed in HEK293T cells. Plasmids encoding human codon optimized MEDLE2 with the N terminal signal peptide removed (aa 2-20) (WT) or encoding the desired point mutations (denoted residues were replaced with alanines) were transfected into HEK233T cells. After 24h, whole cell lysates were prepared, and proteins separated by for SDS PAGE for Western Blot analysis. MEDLE2 mutants were indistinguishable in size from WT MEDLE2. Red, Neomycin; green, HA.

**Figure 6- figure supplement 2.**
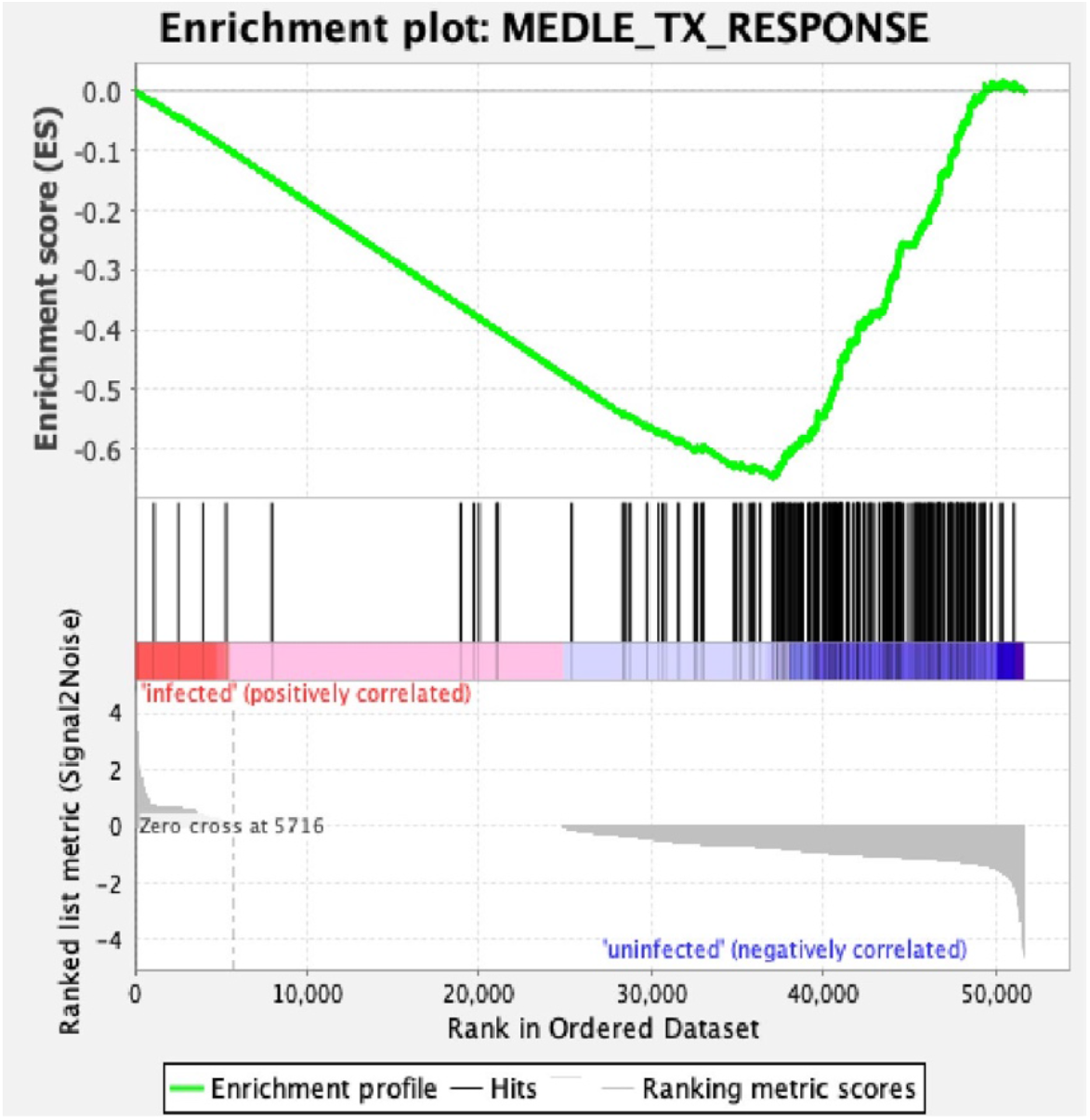
MEDLE2 response signature is absent from intestinal epithelial cells infected with human Rotavirus. Genes with the greatest differential expression (p <0.01, log fold change absolute value > 1.5) were used to define a MEDLE2 gene set from the MEDLE2-GFP transfected cells. RNA sequencing from small intestinal enteroid cultures infected with human rotavirus shows an absence of the MEDLE2 signature created from the MEDLE2 transfection dataset, evidenced by the negative enrichment score resulting from GSEA.

